# DeepWAS: Multivariate genotype-phenotype associations by directly integrating regulatory information using deep learning

**DOI:** 10.1101/069096

**Authors:** Janine Arloth, Gökcen Eraslan, Till F.M. Andlauer, Jade Martins, Stella Iurato, Brigitte Kühnel, Melanie Waldenberger, Josef Frank, Ralf Gold, Bernhard Hemmer, Felix Luessi, Sandra Nischwitz, Friedemann Paul, Heinz Wiendl, Christian Gieger, Stefanie Heilmann-Heimbach, Tim Kacprowski, Matthias Laudes, Thomas Meitinger, Annette Peters, Rajesh Rawal, Konstantin Strauch, Susanne Lucae, Bertram Müller-Myhsok, Marcella Rietschel, Fabian J. Theis, Elisabeth B. Binder, Nikola S. Mueller

## Abstract

Genome-wide association studies (GWAS) identify genetic variants associated with quantitative traits or disease. Thus, GWAS never directly link variants to regulatory mechanisms, which, in turn, are typically inferred during *post-hoc* analyses. In parallel, a recent deep learning-based method allows for prediction of regulatory effects per variant on currently up to 1,000 cell type-specific chromatin features. We here describe “DeepWAS”, a new approach that directly integrates predictions of these regulatory effects of single variants into a multivariate GWAS setting. As a result, single variants associated with a trait or disease are, by design, coupled to their impact on a chromatin feature in a cell type. Up to 40,000 regulatory single-nucleotide polymorphisms (SNPs) were associated with multiple sclerosis (MS, 4,888 cases and 10,395 controls), major depressive disorder (MDD, 1,475 cases and 2,144 controls), and height (5,974 individuals) to each identify 43-61 regulatory SNPs, called deepSNPs, which are shown to reach at least nominal significance in large GWAS. MS- and height-specific deepSNPs resided in active chromatin and introns, whereas MDD-specific deepSNPs located mostly to intragenic regions and repressive chromatin states. We found deepSNPs to be enriched in public or cohort-matched expression and methylation quantitative trait loci and demonstrate the potential of the DeepWAS method to directly generate testable functional hypotheses based on genotype data alone. DeepWAS is an innovative GWAS approach with the power to identify individual SNPs in non-coding regions with gene regulatory capacity with a joint contribution to disease risk. DeepWAS is available at https://github.com/cellmapslab/DeepWAS.

## Background

Genome-wide association studies (GWAS) have been highly successful in identifying genetic variants associated with risk for common diseases and traits ^1^. However, going from pure association to mechanistic insight has been a much more challenging task. Identification of the true causal variants in the context of a disease or trait from within regions of associated variants is hampered by linkage disequilibrium (LD), as nearby variants are likely to co-segregate in a population. The functional variants can, in most cases, not be easily discerned within long stretches of such correlated DNA, which can span several genes and include hundreds of associated variants.

Additional post-processing approaches, so called functional GWAS, have been introduced to provide missing functional annotation to classical GWAS^2^. The majority of published functional GWASs are based on the positional overlap of single-nucleotide polymorphisms (SNPs) with *cis*-regulatory elements such as promoters and enhancers (see Tak and Farnham for a comprehensive review). These functional GWASs indicate that, for common diseases, the majority of associated SNPs reside in non-coding, regulatory regions ^2,4,5^. One drawback of these methods is that the actual impact of each variant on regulatory elements is not assessed, as the annotation is based on positional overlap only. For example, two SNPs that localize to the same chromatin immunoprecipitation with a massively parallel sequencing (ChIP-seq) peak of a transcription factor (TF) ^6^ might have the same, opposing or no functional effects at all. To try to resolve this shortcoming, *in silico* approaches predicting the degree of disruption of TF binding motifs have been used ^7,8^. However, our understanding of the determinants of TF binding to known sequence motifs is still incomplete, limiting the success of such methods. Another post-processing approach to infer functional variants has been annotations that correlated SNP genotype with gene expression or DNA methylation data in the form of expression and methylation quantitative trait locus (eQTL and meQTL) analyses ^9,10^. Additional methods such as binding QTL (bQTL) studies for TF binding ^11^ or massively parallel reporter assays (MPRAs) now add experimental regulatory information on even a single variant resolution^12–14^, yet need to be performed in the cell type of interest. While these approaches can indicate regulatory effects of associated SNPs, they cannot identify single functional variants within an LD block.

In classical, hypothesis-free GWAS, all variants are treated equally despite it being highly unlikely that they are all functionally relevant in every disease-associated tissue of the body. Functional and disease relevant variants may show an association strength below the significance threshold, increasing the efforts required for identifying such regulatory variants. The integration of functional knowledge regarding SNPs into GWAS has the potential to prioritize relevant variants and overcome the current hurdle in gaining mechanistic insights into effects of SNPs associated with a specific phenotype.

Recent advances in systems genetics that harness the predictive power of deep learning might enhance the performance of functional SNP prioritization methods. The deep learning method “DeepSEA” uses only DNA sequence information to predict effects on regulatory chromatin features, such as histone marks, TF binding, or the presence of open chromatin ^15^. For this annotation method, experimental, publicly available data from the ENCODE project ^16^ and the Roadmap Epigenomics Project^17^ for cell type-specific TF binding, histone modifications, and chromatin states were used. This type of functional sequence annotation is superior to *post-hoc*, positional overlap-based methods, as it computes allele-specific differences in the effects of variants on regulatory elements and thus discerns SNPs with predicted functional impact in a given cell type from those just residing within an annotated element by chance. Importantly, the method allows for incorporating cell-type specific regulatory effects of variants at baseline as well as under experimentally challenged conditions, adding additional critical layers to understanding tissue- and context-specific disease mechanisms. This approach was further enhanced to predict tissue-specific gene expression levels and to prioritize putative causal variants associated with human traits and diseases ^18^. Recently, deep learning was successfully applied to predict the impact of non-coding mutations in a family with autism spectrum disorder ^19^ and the clinical impact of single human mutations ^20^. However, in all studies published to date, such deep learning-based annotations have only been applied *post-hoc* to association results from classical GWAS.

We present a conceptually new approach fusing classical and functional GWAS. We obtain regulatory information on SNPs by generating sets of SNPs mapped to so called “functional units” (FUs) using deep learning and then perform multivariate FU-*W*ide *A*ssociation *S*tudies (DeepWAS). A FU is defined as the combination of a specific chromatin feature, a cell type, and an experimental condition. Regulatory SNPs are then grouped based on their predicted capacity to modulate a FU and are jointly subjected to multivariate association with a trait or disease using feature selection with LASSO (L1) penalization^21^. The rationale of this grouping by FUs stems from the idea that as the majority of our chromatin features are TFs (85%, see **Supplementary Table 1**) and that one TF influences a specific cellular function through accumulation of its downstream effects via multiple SNPs on a series of loci ^22^. This optimized variant selection improves the power to identify sets of functional SNPs that may play a role in the etiology of the disease, and map them directly to a specific context – i.e., cell type, transcription factor, stimulation condition. Multiple regression models are relatively new in the field of GWAS, but have already shown promising results ^23–25^. Of note, one of these multi-SNP approaches used pre-clustering of LD blocks prior to LASSO modeling ^26^.

For a proof of concept of this method, we used data from published GWASs of two common diseases, multiple sclerosis (MS) ^27^ and major depressive disorder (MDD) ^28^, as well as the quantitative trait of height ^29^. The heritability of MS and MDD was estimated to be 64% and 40%, respectively ^30,31^, comparable with other common diseases. The heritable nature of height is estimated to be between 70-90% ^32^. GWAS meta-analyses have already identified 180 genetic loci for height ^33^, 200 for MS ^34^, and 44 independent loci for MDD ^35^. We compare the results of DeepWAS in smaller samples to the results from GWAS meta-analyses of the same phenotypes. This allows to identify disease- or trait-associated FUs, generating novel supportive evidence for prior knowledge on pathophysiology. We also complement deepSNPs with QTL networks and generate novel testable hypotheses of disease mechanisms.

## Results

### Directly integrating regulatory information into genotype-phenotype associations (DeepWAS approach)

In order to integrate regulatory information into the classical GWAS approach, knowledge of regulatory information on a single SNP level is required. To retrieve a set of regulatory SNPs with effects on cell type-specific chromatin features, we employ DeepSEA variant effect predictions. We generated reference and alternative allele sequences of 1,000 bp regions centered around each GWAS SNP position (eight million SNPs) and used a pre-trained deep neural network ^36^ to obtain allele-specific probabilities for a genomic feature to manifest in a specific cell type under a given experimental condition (a pair of one chromatin feature in one cell type is further called “functional unit” (FU)). For this study, we used 919 pre-defined FUs (see **Supplementary Table 1**). For example, the FU “p53-baseline-HeLa” was informed by all ChIP-seq peaks of the TF p53 in baseline conditions in the HeLa cell line. We then employed the DeepSEA e-value metric that estimates the impact of a variant on the functional readout by comparing the allele-specific probabilities per variant to one million random SNPs from the 1,000 genomes project ^37^. Only SNPs with significant e-values (e-value<5×10^−5^) were thus selected as likely to be regulatory. This filtering identified 40,000 predicted regulatory SNPs, i.e., a SNP with effects in one FU. We next merged all variants that are predicted to moderate the same FU into SNP sets. To associate these regulatory SNPs with a disease or trait in a multivariate manner, we employ L1-regularized (LASSO) linear regression models including multiple SNPs using the stability selection method. From all 919 FU models, we extracted those regulatory SNPs that show significant trait association and defined them as deepSNPs (see **Methods** and **Figure 1**). Moreover, the DeepWAS approach identifies SNP-phenotype associations directly in a cell-type specific regulatory context.

**Figure 1:**
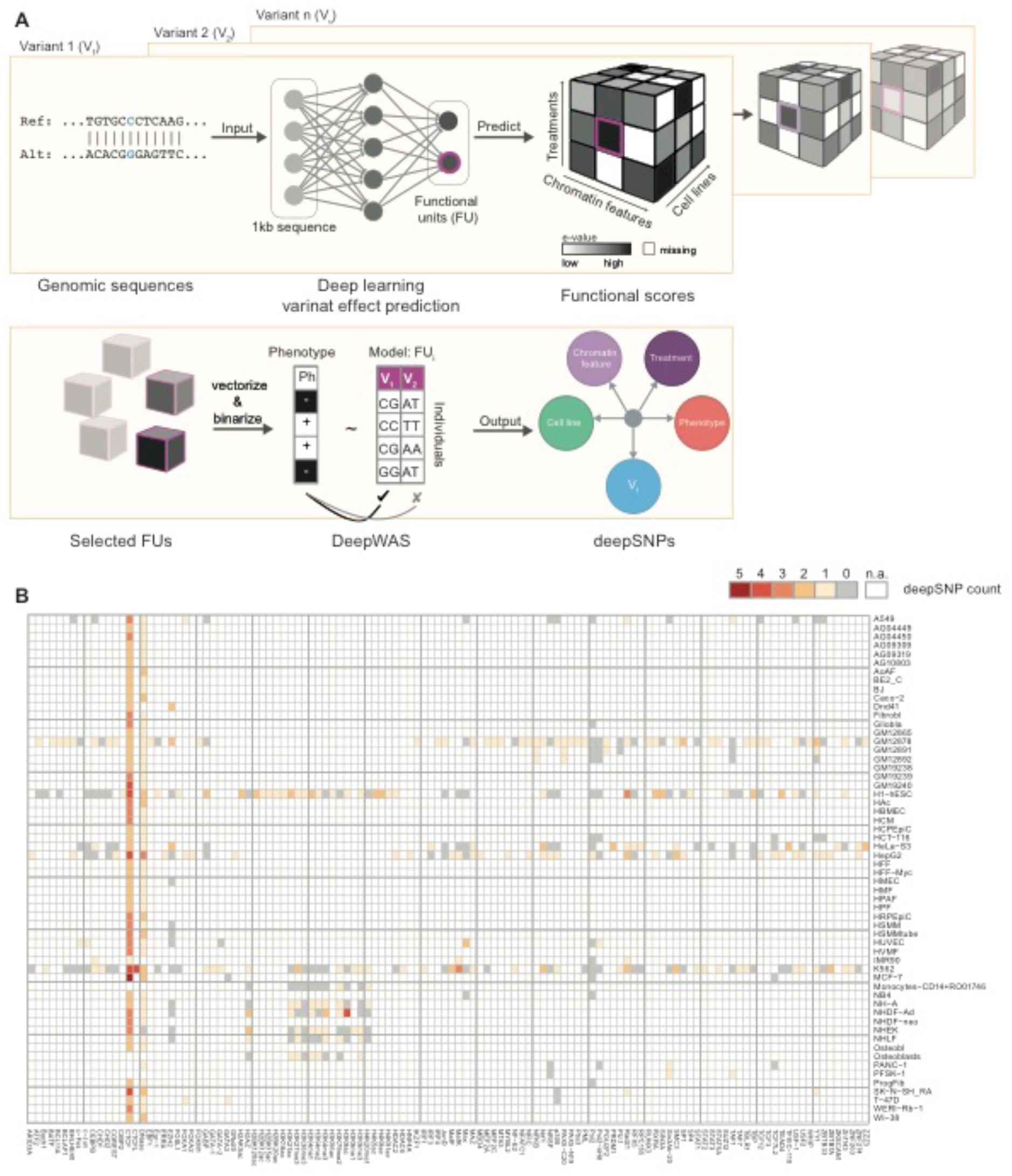
Workflow of DeepWAS. **A)** A deep-learning based framework predicts combined binding probabilities for chromatin features, cell lines, and treatments, called functional units (FU) for 1,000 bp centered around a SNP. FUs are selected for a potential functional role of a variant using a cutoff for functional scores. This process is repeated for all genotyped variants. The genotype-phenotype association is analyzed for each FU using LASSO regression with stability selection. Unlike GWAS, DeepWAS implicates a regulatory mechanism underlying the phenotype of interest with information on relevant cell lines and TFs. **B)** DeepWAS was applied to 36,409 regulatory SNPs that were retained after filtering for allele-specific effects in any given FU. These SNPs were tested for an association with multiple sclerosis (MS). The heatmap shows the number of selected chromatin features vs. cell lines. Chromatin features are limited to be present in at least two distinct cell lines. Missing values, represented in white, show FUs for which no data were available.

To test this approach, we applied DeepWAS to a dataset from an adequately-powered GWAS for MS (KKNMS GWAS, 15,283 individuals from two independent MS case-control cohorts ^27^) and found that for 637 out of 919 FU models, associated with at least one variant (see **Figure 1B**). Most models identified one deepSNP per FU while 147 models identified two, three or four deepSNPs to jointly moderate a FU. In total 53 unique MS-specific deepSNPs, excluding the region encoding the major histocompatibility complex (MHC) on chromosome 6, were identified by DeepWAS. These deepSNPs moderate 120 chromatin features in 133 cell lines (see **Supplementary Table 2**). Note that 16 out of the 53 deepSNPs were in pairwise LD with each other (*r*^2^≥0.5). 111 MS-specific deepSNPs were located in the MHC region. In addition to MS, we analyzed underpowered GWAS datasets for MDD (3,627 individuals recruited for recurrent MDD ^28,38^) and height (3,658 individuals of the population-based KORA cohort ^39^). Sixty-one MDD-specific deepSNPs in 237 FUs (**Supplementary Figure 1A** and **Supplementary Table 3**) and 43 height-specific deepSNPs in 381 FUs (**Supplementary Figure 1B** and **Supplementary Table 4**) were identified in these DeepWAS.

### Characterization of deepSNPs

We followed these results up by further characterizing the identified deepSNPs from the three independent analyses. DeepSNPs for MS and height were more likely to be located in intronic regions (32-33% deepSNPs in first or other introns) while deepSNPs for MDD were more likely to cluster in distal intergenic regions (>3 kb=53% vs. >3 kb=36-37% for MS and height, respectively). MS- and MDD-specific deepSNPs were never found within coding regions (**Figure 2A**). DeepSNPs were always identified in a cell-type specific manner. We thus interrogated the cell type specificity on the level of tissue category, in order to reduce complexity (**Figure 2B**). Note, that tissue categories were influenced by the richness of investigated cell types per category. For example, the tissue group blood encompasses 79 cell types, while brain contained only 14 tissues. Overall, more tissue groups were identified when the number of identified deepSNPs increased. Interestingly, compared to MS and MDD, a lower number of identified deepSNPs for height were relevant in brain tissues. At the same time, a larger number of deepSNPs were active in pancreatic tissue. Notably, an association of height with pancreatic cancer has previously been shown ^40^.

**Figure 2:**
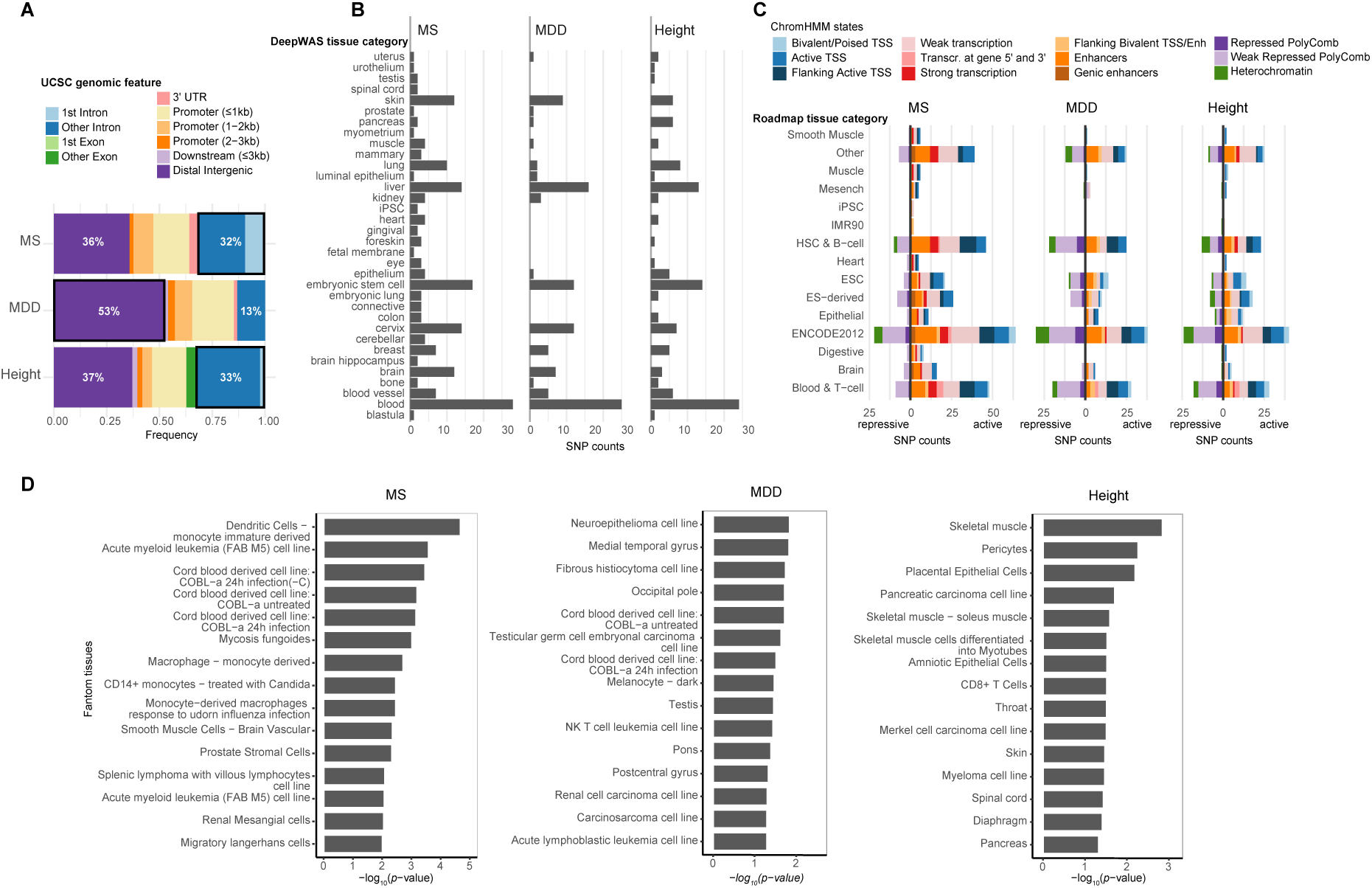
Functional characterization of DeepWAS hits. **A)** Annotation of the genomic regions in which deepSNPs are located: 63-87% of the genomic positions of deepSNPs overlapped with non-coding DNA elements. Seventeen of 53 MS-specific (32%), 14 of 43 height-specific (33%) and 8 of 61 MDD-specific (13%) deepSNPs mapped to introns (first and other introns). Over a half of the MDD-specific deepSNPs (53%) resided in distal intergenic regions (>3 kb). None of the MS- and MDD-specific deepSNPs were located in exons. **B)** Bar plots for each phenotype showing the number of unique deepSNPs annotated to a top-level tissue category (ENCODE). **C)** Overlap of MS-, MDD-, and height-specific deepSNPs with ChromHMM states from Roadmap epigenomes based on top-level tissue group matching. Most of our MS- and height-specific deepSNPs mapped to predicted active chromatin states (82-86%), whereas nearly half of MDD-specific deepSNPs mapped to inactive chromatin states (43%). **D)** Tissue enrichment with FANTOM gene expression data. The top 15 significantly enriched tissues are shown (all *p*-values≤0.05).

As DeepWAS includes only a limited number of histone marks, we next overlapped deepSNPs with predicted chromatin states from the 15-state ChromHMM model^41^ (**Figure 2C**). For both DeepWAS cell types (ENCODE) and 111 epigenomes, we used top-level tissue categories to overlap deepSNPs with chromatin state predictions in the respective matched tissue. We observed tissue and context (disease or trait) specific roles of chromatin states. Most of the MS-specific deepSNPs mapped to active chromatin states (82%, see **Figure 2C**). For MDD-specific deepSNPs a larger fraction overlapped with repressive marks (43%, see **Figure 2C**).

To investigate the tissue specificity of the genomic loci around deepSNPs and to extend the number of disease-relevant tissues, we further tested if our deepSNPs and their proxies (*r*^2^≥0.5) were enriched in the loci of tissue-specifically expressed genes by leveraging Fantom CAGE data ^42^. MS-specific deepSNPs were significantly enriched in the regions active in different immune cell types, height-specific deepSNPs in skeletal muscle cells, pericytes, and also pancreatic carcinoma cell line, and MDD-specific deepSNPs in neurocytoma as well as in different brain regions and immune cells (*p*-value≤0.05, see **Figure 2D**).

### DeepWAS identifies GWAS loci

To evaluate the convergences of DeepWAS and GWAS, we evaluated how many deepSNPs map to cohort-specific results from classical GWAS and to results of larger published meta-analysis of GWAS (**Figure 3A**). In the published GWAS of the KKNMS dataset used in the present study, the MHC region as well as variants at 15 loci outside of this region were significantly associated with MS on a genome-wide level ^27^. When comparing KKNMS GWAS and DeepWAS results on a single SNP level, eleven of the 53 MS-specific deepSNPs or their LD proxies (*r*^2^≥0.5) mapped to eight independent loci (*CLEC16A, MYO15A, PAGR1, CD58, GFI1, EVI5*, and intergenic regions on chromosome 1 and 10). The remaining deepSNPs (n=42) showed an association strength with at least nominal significance in the original GWAS with *p*-values all ≤5.13×10^−4^, but did not reach genome-wide significance. We next overlapped the deepSNPs with results from the GWAS by the International Multiple Sclerosis Genetics Consortium (IMSGC ^43^) that included over 47,000 MS cases and 68,000 controls. Here, a total of 12 deepSNPs mapped to nine independent genome-wide significant loci. Eight of these 12 deepSNPs, corresponding to 8 out of 9 loci, were also genome-wide significant in the KKNMS GWAS^27^.

**Figure 3:**
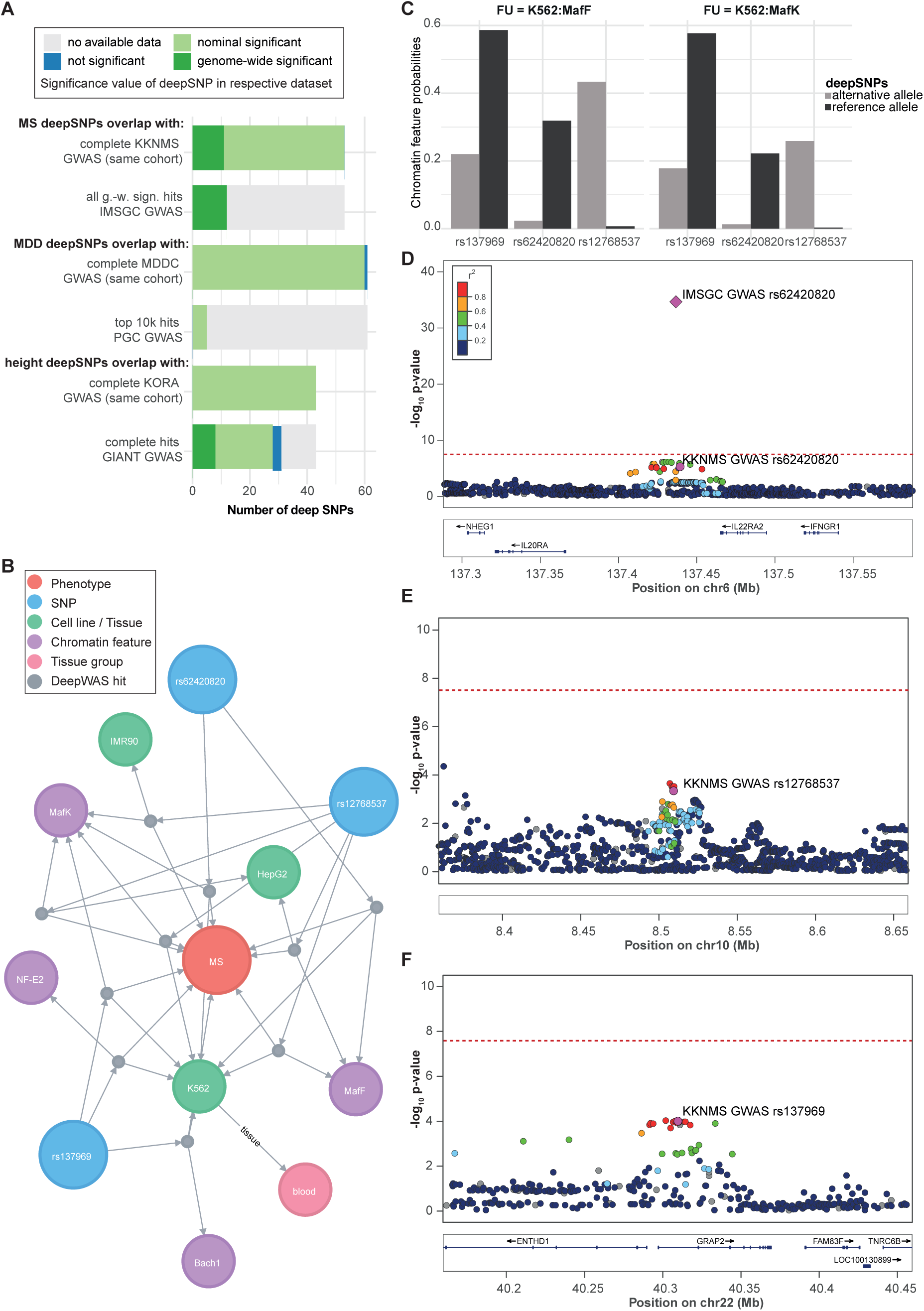
Comparison of DeepWAS vs. GWAS results. **A)** Bar plot of the overlap of cohort-matched GWAS and consortia GWAS SNPs with deepSNPs. g.-w. s=genome-wide significant. **B)** Network of MS-specific deepSNPs generated by using a graph database and showing the deepSNP rs62420820 in the K562 cell line, a genome-wide significant signal in the IMSGC MS GWAS, but sub-threshold in the cohort-specific KKNMS GWAS. Edges represent the association relation of deepSNPs, chromatin features with or without treatment, cell lines, and top-level tissue group. **C)** Bar plots showing the predicted DeepSEA probabilities for deepSNP sequences carrying the alternative and reference allele group by their FU. **D-F**) Locus-specific Manhattan plots of the MS-specific deepSNPs rs62420820, rs12768537, and rs137969, based on classical GWAS. Plots were produced using LocusZoom with EUR samples of the 1,000 genomes November 2014 reference panel on the hg19 build ^69^. Dots represent KKNMS GWAS *p*-values and the diamond shows the IMSGC GWAS signal *p*-value. Color of the dots indicates LD with the lead variant = deepSNP (magenta), grey dots have LD *r*^2^ missing.

For MDD, 60 out of the 61 MDD-specific deepSNPs or their proxies (*r*^2^≥0.5) reached nominal significance in a univariate GWAS from the same cohort (MDDC n=3,627 individuals, no genome-wide significant association). Five MDD-specific deepSNPs overlapped with the top 10,000 SNPs of the recently published largest GWAS for MDD by the PGC ^35^ (included 135,458 MDD cases and 344,901 controls; PGC GWAS deepSNP max. *p*-value ≤ 3.1 ×10^−4^), which represents a 2.2 fold enrichment (permutation *p*-value=0.071) over randomly sampled regulatory SNP sets (*n*=31,929 regulatory SNPs) and their LD proxies. These five deepSNPs map to five independent loci: *ZFPM2-AS1, SBK2, MTFR1*, and two intergenic regions on chromosome 7 (nearby genes: *WNT2* and *ASZ1*) and a locus on chromosome 8 (nearby genes: *LINC00293* and *LOC100287846*).

All of the 43 height-specific deepSNPs reached nominal significance in the classical GWAS in the same cohort (KORA cohort n=3,658 individuals, no genome-wide significant association) with a max. *p*-value ≤7.7×10^−3^. Eight of the 43 height-specific deepSNPs mapped to seven independent genome-wide significant loci (*DIS3L2, ZBTB38, LCORL, PDLIM4, ZNF311, HABP4*, and *PXMP4*) of the latest GWAS from the GIANT Consortium, which included over 183,727 individuals ^33^.

In all the three DeepWAS, this approach identified regulatory SNPs that overlapped not only with SNPs associated with the same traits in the cohort-matched GWASs, but also with genome-wide significant associations from the larger consortia GWASs for these traits. These associations had not survived correction for multiple testing in the underpowered cohort-matched GWAS, but were detected using DeepWAS.

### Deriving hypotheses on disease-associated mechanisms in MS from DeepWAS result

We next wanted to illustrate how DeepWAS can accelerate the discovery of disease mechanisms. Within the DeepWAS results for MS, we identified, for example, the intergenic region *IL20RA-IL22RA2* on chromosome 6 that includes deepSNP rs62420820 (**Figure 3B**), which was genome-wide significant in the IMSGC GWAS (*p*-value=9.26×10^−36^, **Figure 3D**) and nominally significant in the KKNMS GWAS conducted on the MS cohort used for the DeepWAS analysis (*p*-value=1.23×10^−5^, **Figure 3D**). In comparison to the published GWAS-based results, DeepWAS adds the novel and testable hypothesis that the TFs MafF and MafK contribute to MS susceptibility. The IMSGC GWAS and deepSNP rs62420820 shows allele-specific TF binding differences for MafF and MafK in the leukemia cell line K562 (**Figure 3C**). Of note, additional deepSNPs were identified within the FUs MafK:K562 (**Figure 3B/E**: deepSNP rs12768537 on chromosome 10 and **Figure 3B/F**: deepSNP rs137969 on chromosome 22) and MafF:K562 (deepSNP rs12768537), supporting a role of these TFs in the etiology of MS.

We also identified deepSNPs that were detected at a genome-wide significance level by both the cohort-matched KKNMS GWAS and IMSGC GWAS. This includes deepSNP rs1985372 on chromosome 16, located in the *CLEC16A* locus, previously suggested as a candidate gene for MS (the deepSNP rs1985372 was significant in the KKNMS GWAS on the same cohort and is in complete LD with the SNP rs2286974 (*r*^2^=0.99), which was significant in the IMSGC GWAS ^27,43^). DeepWAS now adds the regulatory information that these SNPs alter TF binding of GABP, GATA-1, GATA-2, p300, STAT1, STAT2, STAT5A, and TBLR1, all expressed in K562 cells, and that these TFs play a MS-specific role in regulation of *CLEC16A*.

### Regulatory effect of MS-specific deepSNP loci

Allele-specific effects on chromatin features and TF binding are likely to be reflected by changes in DNA methylation and gene expression. To test whether the MS-specific deepSNPs or their proxies (*r*^2^≥0.5) associate with differences in gene expression and DNA methylation, we used publicly available *cis*-meQTL, *cis*-eQTL, and eQTM data from multiple resources: 1) The Biobank-Based Integrative Omics Study (BIOS) analyzing whole blood (>2000 samples) ^44,45^, the CommonMind Consortium (CMC ^46^) dorsolateral prefrontal cortex data (DLPFC) (n=603 samples), and GTEx data ^47^. In the largest resource, BIOS, we observed that 36 of the 53 non-MHC, MS-specific deepSNPs were significant meQTLs (68%) and 20 significant eQTLs (38%), which represent a 1.7 and 1.9 fold enrichment over randomly sampled regulatory SNPs and their assigned LD proxies, respectively (permutation *p*-values≤0.011, see **Figure 4A** and **Supplementary Table 5)**. We next restricted DeepWAS hits to deepSNPs moderating immune cell lines (n=13 deepSNPs, 46 chromatin features, 4 cell lines and 50 FUs, **Supplementary Table 2**), given the relevance of this tissue in MS ^48^. Of this subset of 13 immune MS-specific deepSNPs, 62% (n=8 deepSNPs) overlapped with meQTL variants (meSNP) and 38% (n=5 deepSNPs) with eQTL variants (eSNP) in BIOS. In their recent study, the IMSGC identified significant eQTL effects in naive CD4+ T cells and monocytes for only 18% of their significant GWAS loci ^49^. When overlapping our immune MS-specific deepSNPs with the same QTL datasets, we found 43% (n=23 SNPs) of these deepSNPs to be part of eQTLs in CD4+ T cells and 38% (n=20 deepSNPs) in monocytes. In addition to publicly available data, we also conducted an eQTL analysis using blood gene expression levels from a subset of 319 MS patients of the MS cohort. In total, 47% (n=25 deepSNPs) of our MS-specific deepSNPs showed a significant eQTL effect in this data (**Figure 4A** and **Supplementary Table 6**), of which 14 deepSNPs had previously been identified as part of eQTLs in blood. In the GTEx database, we found 42% of these deepSNPs to have a significant effect in blood eQTLs (**Supplementary Table 8**).

**Figure 4:**
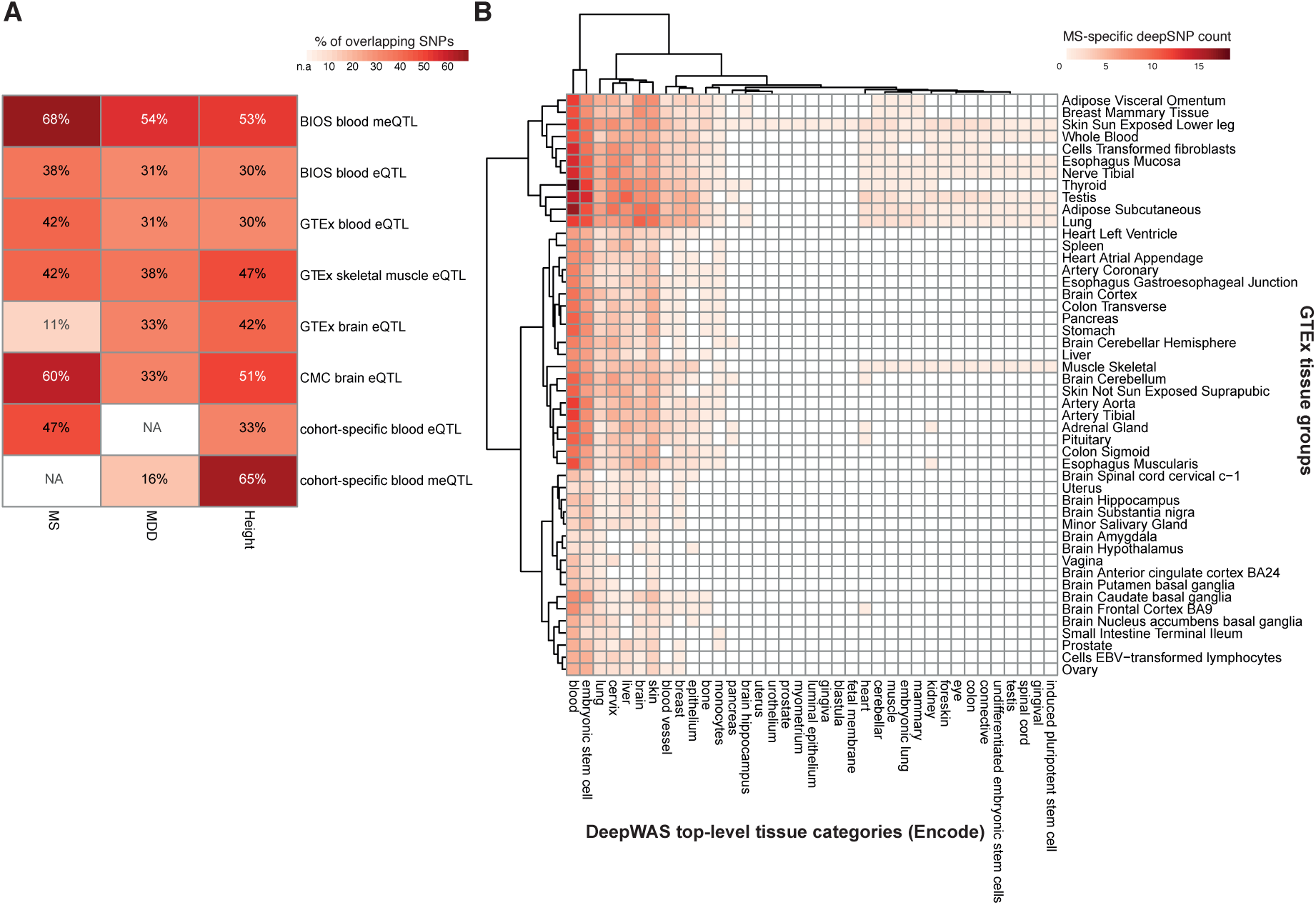
Context-related regulatory capacity of deepSNPs. **A)** Heatmap showing the percentage of overlap of MS-, MDD-, and height-specific deepSNPs or their proxies (*r*^2^≥0.5) with *cis*-meQTL and *cis*-eQTL data from multiple resources, see also **Supplementary Tables 5-16. B**). Heatmap depicting GTEx tissue groups and DeepWAS top-level tissue category overlap among the MS-specific deepSNP FUs.

Taken together, QTL data from various resources support that the DeepWAS of MS identified SNPs in transcriptionally relevant regions and emphasize their putative regulatory role in immune function.

### Regulatory effect of MDD and height deepSNP loci

The 61 MDD- and 43 height-specific deepSNPs were also transcriptionally active in the respective relevant tissues (see **Figure 4A** and **Supplementary Tables 9-16**) and tagged more BIOS eQTLs and meQTLs than expected by randomly sampling of regulatory SNPs and assigning their LD proxies (MDD: fold enrichment for eQTLs=1.7 and meQTLs=1.5, permutation p-value=0.009 and 0.015, respectively; height: fold enrichment for eQTL=1.5 and meQTLs=1.5, permutation p-value=0.068 and 0.013, respectively, see Methods). Interestingly, in the much larger PGC GWAS only 20% of their 44 genome-wide significant hits overlapped with BIOS meQTLs ^35^, while the overlap was 54% for MDD-specific deepSNPs (**Figure 4A**). For a subset of both the MDDC and the KORA cohort, methylation levels and/or expression levels were measured. We therefore calculated meQTL and eQTL effects and found 16% of MDD-specific deepSNPs with significant meQTL effects, 65% of the height-specific deepSNPs with meQTL and 33% with eQTL effects (see **Figure 4A** and **Supplementary Tables 10,14-15**).

All data from DeepWAS results with additional QTL information is stored in a searchable graph database and accessible at https://github.com/cellmapslab/DeepWAS.

### Disease mechanisms on the level of functional units

While DeepWAS can be used to predict the phenotype from the genotype, it is also interesting to annotate the relationship of FUs to disease or trait. For example, MS is an immune-mediated disorder affecting the central nervous system (CNS). Naturally, the CNS is difficult to directly examine as a study tissue. DeepWAS might here be used to identify a good proxy study tissue. To interrogate this hypothesis, we counted the number of MS-specific deepSNPs of each MS associated FU tissue group that overlapped with eSNPs of a GTEx tissue group (**Figure 5B** and **Supplementary Figure 2**). The MS heatmap in **Figure 4B** support the hypothesis that a proxy tissue can be identified and suggests that blood deepSNPs are surrogates for brain tissues and other tissues, especially for the thyroid.

**Figure 5.**
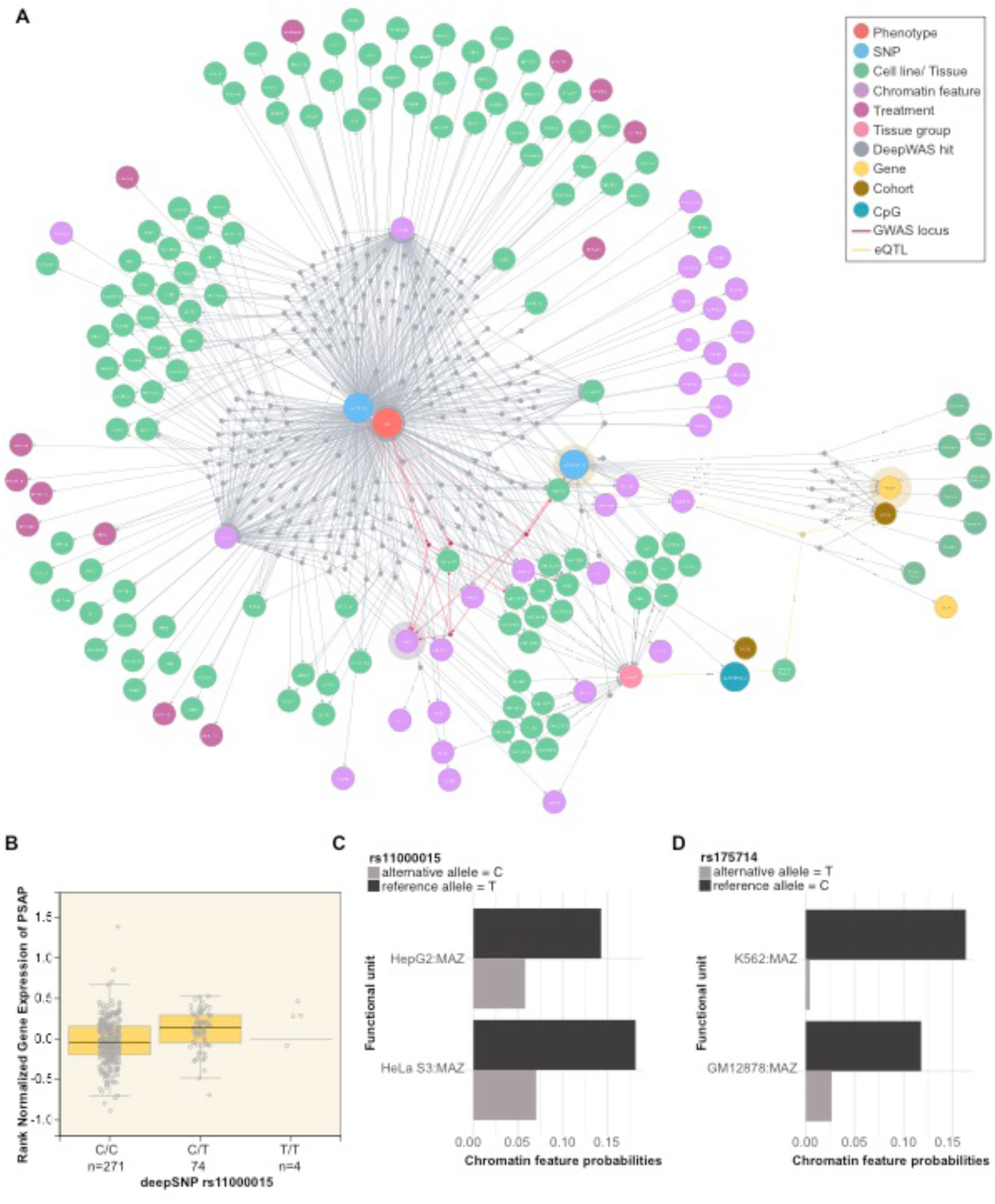
QTL network. **A)** Network showing one of the putative key regulators for MS, deepSNP rs175714 on chromosome 14. DeepSNP rs175714 is associated with differential TF binding of the TF MAZ, one of the top-associated loci in the KKNMS GWAS, where no significant transcriptional effect could be identified in the *post-hoc* analysis. Edges represent the associations between deepSNPs and chromatin features with or without treatment, cell lines, top-level tissue group, CpGs, and genes through dummy nodes identified either using DeepWAS or QTLs. Dummy nodes are used for preserving all entities of deepSNP and QTL associations. Edges highlighted in red show the DeepWAS results for MAZ, in yellow show the eQTL connections illustrated in B, and shades refer to downstream QTL results shown in B. **B)** Box plot of GTEx whole blood eQTL data showing the relationship between *PSAP* gene expression and deepSNP rs11000015 genotype. **C-D**) Chromatin feature probabilities for the significant FU of the deepSNP sequences carrying the reference (black) and alternative (gray) allele.

Moreover, DeepWAS results can be used to identify single SNPs as key regulators, i.e., those SNPs with effects on multiple chromatin features. DeepWAS identified, for example, the intergenic deepSNP rs175714 on chromosome 14 as a key regulator for MS (**Figure 5A-B**). It affects the binding of multiple chromatin features at the same time (n=29) in 116 cell types (**Figure 5A-B**). One of these chromatin features is the TF MAZ. MAZ itself is one of the top-associated loci of the KKNMS GWAS (tag SNP rs34286592 on chromosome 16, *p*-value=4.58×10^−10^), but no significant transcriptional effect was previously identified in a *post-hoc* analysis of the GWAS ^27^. Interestingly, the MS-specific deepSNP rs175714, together with the MS-specific deepSNP rs11000015 on chromosome 10, had a significant effect on the binding of MAZ and they were jointly associated with MS disease status. The deepSNP rs11000015 is correlated with expression levels of the Prosaposin (*PSAP*) gene in multiple tissues, whole blood gene expression levels of PSAP are shown in **Figure 5C**.

Another example is the TF MEF2C in the analysis of MDD, where DeepWAS identified the intergenic SNP rs7839671 on chromosome 8 as one of the key regulators for MDD (see **Supplementary Figure 3**). *MEF2C* is an important risk gene for MDD and the *MEF2C* gene itself is one of top-associated loci of the PGC GWAS for MDD ^35^ (*MEF2C*, chr5: 87.443-88.244 Mb, p= 7.9×10^−11^, **Supplementary Figure 3**). SNP rs7839671 and its proxies are associated with differences in mRNA expression of *SPIDR* and *MCM4* and show meQTLs with an intergenic region (for more details, see **Supplementary Figure 3**).

### QTL network analyses

In-depth investigation of the wealth of additional regulatory capacities of deepSNPs were carried out by generating QTL-networks that combine all pairwise links of meQTL (SNP-CpG), eQTL (SNP-gene), eQTM (CpG-gene), and deepSNP-FU information. QTL network analyses helped us to identify the SNPs that showed joint effects on epigenetic and transcriptomic levels, i.e., deepSNP = eSNP = meSNP where the deepSNP harbors an eQTM. The resulting networks are called three-way QTL interaction networks. Three MS-specific deepSNPs on chromosome 17, rs2273030, rs4925172 (both in complete LD with each other *r*^2^=1), and rs7207666 (*r*^2^=0.7) are located in the *SHMT1* locus, a candidate gene for MS ^27,49^ (**Supplementary Figure 4A**). The genetic variants and *SHMT1* are connected via eQTL, meQTL, and eQTM relations (**Supplementary Figure 4B**). In comparison to the previously published findings on the locus, DeepWAS informs that the TF Yin Yang 1 (YY1), expressed in multiple cell lines, may lead to a downregulation of *SHMT1* gene expression and hypermethylation of cg25492364 and cg26763362, thus connecting the disease associated SNPs with specific regulatory functions.

Moreover, a three-way QTL interaction network analysis identified the height-specific deepSNP rs7146599 on chromosome 14 to affect a cascade of eight chromatin features in eleven cell lines (see **Supplementary Figure 5A-B**). The network also included rs2871960 on chromosome 3, linked to the *ZBTB38* locus and correlated with multiple CpG sites. *ZBTB38* has been previously shown to play an important regulatory role in height ^50^.

Thee-way QTL interaction network analysis identified the MDD-specific deepSNP rs163105 on chromosome 5 to alter the expression of *SKIV2L2* (also known as *MTR4*). This gene has already been shown to be differentially expressed between depressed women and controls (see **Supplementary Figure 6)**. Furthermore, *SKIV2L2* has been implicated in the stress response and neurodegeneration through the nuclear exosome-targeting (NEXT) complex ^52^.

In summary, DeepWAS allows the direct identification of putative master regulators, TFs and chromatin features for a subsequent in-depth analysis of genetic association signals.

## Discussion

In classical GWASs, all SNPs are tested independently from each other on a genome-wide scale, thereby implicitly assuming that any SNP can moderate the function of any cell state at any time. It is now clear that disease associations, especially regarding common disorders, are driven by SNPs altering the function of regulatory elements. Hence, it is likely that testing all SNPs in GWAS is not necessary but rather we should use functional annotation to help prioritize putative risk variants ^17^. So far, several *post-hoc* functional annotations of GWAS results have been reported ^53–55^. In this manuscript, we present DeepWAS, a novel analysis tool for genetic associations that fuses GWAS and functional annotation into one single step (**Figure 1**). We employ the powerful deep learning based method DeepSEA to predict regulatory effects of chromatin features in various cell types on a single SNP level ^36^. In addition, we implement multi-SNP regression models with L1 regularization to identify so-called deepSNPs. The SNPs of one FU are only selected when they are jointly associated with the disease or trait. To the best of our knowledge, this study is the first to combine deep learning-based predictors with multivariate models. By applying DeepWAS to three datasets, we show that this method allows direct fine-mapping of GWAS associations at a single base resolution as well as direct functional annotation of the association signal to both chromatin features and cell types, generating novel mechanistic hypotheses. We also demonstrate that DeepWAS, by pre-selecting functionally relevant SNPs and integrating multivariate statistics, might increase power to detect true positive signals.

We applied DeepWAS on a well-powered GWAS dataset for MS (n=15,283 individuals), but also to underpowered GWAS data for MDD (n=3,627 individuals) and height (n=3,658 individuals). In all three phenotypes, many SNPs in many genes contribute to genetic variation in the population and the effect size of each SNP was found to be small ^56^. Therefore, large sample sizes are needed to discover additional risk variants. We identified 35 putative new candidate MS risk SNPs outside of the MHC region that did not reach genome-wide significance in GWAS (in total: 53 non-MHC, MS-specific deepSNPs). For MDD, DeepWAS prioritized 56 new putative risk variants (in total: 61 MDD-specific deepSNPs) and for height 35 new risk variants (in total: 43 height-specific deepSNPs), even though the classical GWAS approach for MDD and height did not yield genome-wide significant results. Importantly, when comparing our deepSNPs identified in these smaller cohort-matched GWASs to large consortia GWASs of the same phenotype, DeepWAS was able to detect SNPs that reached genome-wide significance in these large consortia GWASs but not in GWAS of the same sample (**Figure 3**). Importantly, all deepSNPs were associated at least with nominal significance in cohort-matched GWAS, with a subset of deepSNPs reaching genome-wide significance. Interestingly, when comparing our deepSNPs to the larger consortia GWASs, DeepWAS detected SNPs that reached genome-wide significance in these GWASs but not in the smaller-scale GWAS on the individual cohorts (**Figure 3**). For example, 23% of the 53 MS-specific deepSNPs were previously identified in the ISMGC MS GWAS including more than 135,000 individuals (>47,000 MS cases). One-third of these deepSNPs were not detected in the univariate KKNMS GWAS using the same cohorts as in DeepWAS (n>15,000 individuals). The increased power of DeepWAS is underlined by the observation that when, for example, all regulatory MS SNPs (n=36,409 SNPs), without grouping to FUs, were used as input to a single LASSO regression model with stability selection, only 19 SNPs showed a significant association with MS, of which four mapped outside the MHC region. This is in contrast to the 164 MS-specific deepSNPs identified using DeepWAS, of which 53 were outside of the MHC region. Notably, these 36,409 SNPs reside in only 25,000 independent loci, which is also reflected by the fact that 16 out of 53 MS deepSNPs are not independent from each other. The regulatory genotypes have, thus, little correlation and are therefore suitable to be subjected to a L1 penalization. In summary, DeepWAS can increase the power to detect associations of a phenotype with regulatory variants, with genome-wide significant consortia GWAS variants detected using DeepWAS in smaller samples in which they only showed sub-threshold signals using classical GWAS. As outlined below, functional analyses of identified deepSNPs suggest that the additional signals discovered using DeepWAS but not using classical GWAS reside within known, disease-relevant functional pathways and thus likely constitute true-positive associations.

Particularly for non-coding regulatory SNPs, DeepWAS has an advantage over GWAS followed by *post-hoc* annotation and allows to identify transcriptionally relevant regions in the disease context (**Figures 2-5**). In fact, in all three DeepWAS, deepSNPs were identified in cell types and enhancers previously shown to be relevant for the tested phenotype. For example, 47% of the MS-specific deepSNPs (n=35) affect the binding of chromatin features in hematopoetic tissue, and another 30% affect chromatin features in brain tissue or spinal cord (n=16; **Figure 2B**). These findings revealed that the genetic disease risk is driven by altered binding of chromatin features mainly in these two tissues.

DeepWAS results also pointed towards convergent regulatory mechanisms of specific TFs in both MS and MDD. For both disorders, DeepWAS identified a set of SNPs modulating binding of a TF that was found to be associated with the disorders in GWAS. This suggests that SNPs associated with the gene encoding the TF and SNPs altering its binding to target transcripts jointly affect the phenotype. The gene *MAZ* on chromosome 16, for example, has been previously identified as a genome-wide significant GWAS locus for MS ^27^. DeepWAS identified several deepSNPs that include the TF MAZ in a FU (**Figure 5**). DeepSNP rs11000015 on chromosome 10 and deepSNP rs175714 on chromosome 14 alter *MAZ* binding. Supporting evidence for the importance of deepSNP rs11000015 as a risk factor for MS was provided by it being a multi-tissue GTEx eQTL involving *PSAP. PSAP* codes for prosaposin, a precursor of several small nonenzymatic glycoproteins termed sphingolipid activator proteins that assist in the lysosomal hydrolysis of sphingolipids ^57^. Sphingolipids are the main components of nervous tissue, and have been previously linked to MS ^58^. As a second example, we identified MDD-specific deepSNPs altering the binding of TF MEF2C (**Supplementary Figure 3**). SNPs in the locus encoding *MEF2C* are the top signal in the latest meta-analysis for major depression^35^. The MEF2 TF family has been reported to play a major role in synaptic plasticity, which is thought to be disturbed in MDD, especially in the context of stress. Chen and colleagues ^59^ identified the TF MEF2 as a master regulator of developmental metaplasticity, which is important for guiding developmental structural and functional neuronal plasticity. Additional evidence was found by Barbosa *et al*. ^60^, relating MEF2 to activity-dependent dendritic spine growth and suggesting that this TF may suppress memory formation.

To support the validity of the predicted regulatory effects of our DeepWAS associations, we provided multiple lines of evidence that deepSNPs and their surrounding loci are indeed functionally active in their respective tissue. We chose to prioritize DeepWAS results based on additional regulatory impact of DNA methylation and gene expression as shown by their meQTL and eQTL effects. DeepSEA together with eQTL data, has been previously applied to prioritize disease-associated variants ^61^. We observed that 68% of MS-specific deepSNPs are meQTLs and 38% eQTLs in the largest QTL resource, BIOS. When using only random sets of regulatory SNPs with no disease association, no such overlap was found. For all investigated phenotypes, we observed a significant overlap between deepSNPs and meQTL SNPs (**Figure 4**, fold enrichment ≥1.5 and permutation *p*-value over 1,000 random sets ≤ 0.015. For MDD- and MS-specific deepSNPs we found a significant overlap with eQTL SNPs (**Figure 4**, fold enrichment ≥1.5 and permutation *p*-value over 1,000 random sets ≤ 0.011). Moreover, as MS is a disorder developing in the peripheral immune system, we investigated if our deepSNPs alter the expression of CD4+ T cells or influence the expression in monocytes, and found twice as many eQTL effects for deepSNPs as described in the published IMSGC GWAS for MS (43% in CD4+ T cells and 38% in monocytes found using DeepWAS *vs*. 18% in both cell types in the IMSGC GWAS). Using expression profiles from GTEx, we found more MS-specific deepSNPs for blood eQTL SNPs than in brain-related eQTL SNPs (42% *vs*. 11%). This supports the theory that MS is most likely initially triggered by perturbation of immune responses, but that also the functional responses of brain cells are altered and may have a role in targeting an autoimmune process to the CNS.

Finally, we explored whether generating QTL interaction networks of our deepSNPs and extracting the SNPs with an impact on methylation and expression, which also coincidently an eQTM, can help identify likely functional risk mechanisms. We identified deepSNPs in the *SHMT1* gene, a published MS GWAS locus [27,28], where DeepWAS QLT network analysis pinpoints the TF Ying Yang 1 (YY1), expressed in multiple cell lines including immune related cells, as a potential novel risk factor (**Supplementary Figure 4**). YY1 is a ubiquitously expressed TF shown to be essential for B cell development ^62^ and serves as master regulator of T cell exhaustion ^63^. Such dysregulation of immune related cells has been shown to promote MS progression ^64,65^. Additional SNP-protein association studies (pQTLs) ^66^ that show how the deepSNP rs2273030 alter YY1 protein abundances could be extended to develop clinical applications in the context of MS.

DeepWAS is mostly limited by the comprehensiveness of regulatory element catalogues like ENCODE and Roadmap. ENCODE lacks, for example, information for a number of relevant disease-specific stimulation conditions as well as disease-related tissues. We have previously reported on the importance of testing SNPs in stimulated conditions, and shown that glucocorticoid response moderating SNPs only become apparent in the stimulation condition and are not overlapping with baseline eQTLs ^67^. Of note, the glucocorticoid receptor itself is central for the stress response and has been previously implicated in the pathogenesis of MDD ^67,68^. In addition, data from cell lines or bulk tissues will miss variants with effects only specific cell types as well as cell type specific effects dependent on the systemic, developmental and/or tissue context. It is therefore important to be able to retrain the DeepSEA neural network with additional publicly available chromatin features and as well as newly generated experimental data. This will be possible using the DeepWAS pipeline publicly available at https://github.com/cellmapslab/DeepWAS.

In summary, our results indicate that DeepWAS, a method combining deep learning based functional SNP annotation and considering a possible multivariate effect of SNPs to moderate a trait or disease, is a powerful tool to uncover disease mechanisms for common disorders and traits. It also allows direct identification of regulatory SNPs by having a single base resolution and not being limited by the LD structure of the locus, since our regulatory SNPs are mostly independent and we jointly analyze regulatory SNPs only if they are predicted to modulate the same FU (see **Methods**). With ever increasing amounts of available functional data, the DeepWAS approach will become even more valuable and allow integration of both publicly as well as unpublished data generated by individual labs. DeepWAS is a versatile, publicly available tool that can be applied to any GWAS dataset in conjunction with the code available for DeepSEA. While we tested DeepWAS in small and medium size samples, and observed a potential increase in power in detecting phenotype-relevant functional SNPs, applying this method to very large datasets will be even more informative.

## Supporting information

## Supplementary Figures

**Supplementary Figure 1:** The heatmaps showing the number of selected chromatin features vs. cell line for **A**) MDD and **B**) height. Chromatin features are limited to be at least present in two distinct cell lines.

**Supplementary Figure 2:** Heatmaps depicting GTEx tissue groups and DeepWAS top-level tissue category overlap with the **A**) MDD- and **B**) height-specific deepSNPs.

**Supplementary Figure 3: A**) Graph-based QTL Network visualization of the DeepWAS results involving the TF MEF2C, which itself is one of the top associated PGC GWAS loci. Two deepSNPs are jointly associated with MDD and belong the same FU: MEF2C:GM12878. Edges represent the association relation of deepSNPs, chromatin features with or without treatment, cell lines, top-level tissue group, CpGs, and genes. Edges of the FU MEF2C:GM12878 are colored in red. Circular shades mark the corresponding genes or CpGs with are plotted in B and C. **B**) Box plot of GTEx frontal cortex eQTL data showing relationship between *SPIDR* gene expression and deepSNP rs7839671. **C**) Boxplot of recMDD meQTL data illustrating relationship between cg01650371 methylation and rs10099827 genotype in recMDD samples. Variant rs10099827 is a proxy of deepSNP rs7839671 (*r*^2^=0.8). DeepSNP rs7839671 exhibits a meQTL effect on the same CpG and was excluded from the original meQTL analysis, as it is 571 kb away from the CpG site (meQTL distance cutoff≤250 kb).

**Supplementary Figure 4: A**) Locus-specific Manhattan plot of the deepSNP rs2273030 that is a sub-threshold GWAS SNP for MS. The plot is produced using LocusZoom with EUR samples of the 1,000 genomes November 2014 reference panel on the hg19 build ^69^. Dots represent GWAS *p*-values and the color of dots indicates LD with the lead variant, grey dots have LD *r*^2^ missing. **B**) MS-specific three-way QTL interaction network generated by using a graph database and highlighting only the deepSNPs with eQTL and meQTL effects that also harbor an eQTM. Edges represent the association relation of deepSNPs, chromatin features with or without treatment, cell lines, top-level tissue group, CpGs, and genes.

**Supplementary Figure 5: A**) Height-specific three-way QTL interaction network highlighting deepSNP rs7146599 on chromosome 14 as one of the moderators of height in eleven cell lines. It affects a cascade of chromatin features (n=8) and shows meQTL and eQTL effects that, at the same time, harbor an eQTM. The CpG site and the deepSNP thus affect the transcriptional level of the same genes. In addition, the network includes rs2871960 on chromosome 3, linked to the *ZBTB38* locus and correlated with multiple CpG sites. Edges represent the association relation of deepSNPs, chromatin features with or without treatment, cell lines, top-level tissue group, CpGs, and genes. **B**) Heatmap illustrating the deepSNPs affecting the binding of more than four chromatin features.

**Supplementary Figure 6:** MDD-specific three-way QTL interaction network generated by using a graph database and highlighting only the deepSNPs with eQTL and meQTL effects that also harbor an eQTM. It shows that the MDD-specific deepSNPs rs163105 on chromosome 5 changes the expression of *SKIV2L2* (also known as *MTR4*). Edges represent the association relation of deepSNPs, chromatin features with or without treatment, cell lines, top-level tissue group, CpGs, and genes.

**Supplementary Figure 7:** Lasso Stability selection results for FU: MEF2C-GM12878: the y-axis indicates number of boosting iterations, the x-axis indicates the stability selection probability, and the horizontal line correspond the 0.7 probability threshold.

## Supplementary Tables

Supplementary Table 1: List of all functional units

Supplementary Table 2: List of MS-specific deepSNPs

Supplementary Table 3: List of MDD-specific deepSNPs

Supplementary Table 4: List of height-specific deepSNPs

Supplementary Table 5: MS-specific deepSNP and BIOS QTL overlap

Supplementary Table 6: MS-specific deepSNP and MS-patient eQTL overlap

Supplementary Table 7: MS-specific deepSNP and CMC eQTL overlap

Supplementary Table 8: MS-specific deepSNP and GTEx eQTL overlap

Supplementary Table 9: MDD-specific deepSNP and BIOS QTL overlap

Supplementary Table 10: MDD-specific deepSNP and MDD-patient meQTL overlap

Supplementary Table 11: MDD-specific deepSNP and CMC eQTL overlap

Supplementary Table 12: MDD-specific deepSNP and GTEx eQTL overlap

Supplementary Table 13: Height-specific deepSNP and BIOS QTL overlap

Supplementary Table 14: Height-specific deepSNP and KORA meQTL overlap

Supplementary Table 15: Height-specific deepSNP and KORA meQTL overlap

Supplementary Table 16: Height-specific deepSNP and GTEx eQTL overlap

## Methods

### Clinical Samples

#### Major depressive disorder cohorts (MDDC)

Two MDD cohorts, recMDD and BoMa, were analyzed. The recMDD cohort consists of 1,774 Caucasian individuals recruited at the Max-Planck Institute of Psychiatry (MPIP) in Munich, Germany and two satellite hospitals in the Munich metropolitan area (BKH Augsburg and Klinikum Ingolstadt): 756 controls (581 women, 275 men) and 879 cases diagnosed with recurrent major depression (585 women, 294 men). Please see Muglia *et al*. ^28^ for more details on sample recruitment and characterization. The BoMa cohort consists of 1,889 Caucasian individuals: 1,292 controls (656 men, 636 women), 597 (212 men, 385 women) of which had a depressive disorder. Recruitment strategies and further characterization have been described previously ^70^.

#### Multiple sclerosis cohorts (KKNMS)

Two MS cohorts, referred to as DE1 and DE2, were analyzed. Both data sets included patients diagnosed with either MS or the prodromal clinically isolated syndrome. DE1 consists of 3,934 cases and DE2 consists of 954 cases; for more details see Andlauer *et al*.^27^. Controls for these cohorts were obtained from several cohorts across Germany: KORA^39^, HNR ^71^, SHIP ^72^, DOGS ^73^, FoCus ^74^, PopGen ^75^, recMDD ^28^, and MARS ^76^; for more details see Andlauer *et al*. ^27^.

#### Population-based cohort (KORA)

The study population consisted of participants from the KORA (Kooperative Gesundheitsforschung in der Region Augsburg) study ^39^, which has been collecting clinical and genetic data from the general population in the region of Augsburg, Germany for more than 20 years. Here, the independent cohorts S3 and S4, including their follow-ups (F3 and F4), were analyzed.

### Genotype data and imputation

#### Data generation

Genotype data was generated for each cohort individually. Details on the methods used can be found in the individual papers (recMDD ^28^, BoMa ^70^, KKNMS ^27^, and KORA ^39^).

#### Quality control and imputation

Quality control (QC) of KKNMS genotype data and imputation have been previously described ^27^ and the same pipeline was applied for KORA and recMDD genotype data. The QC was conducted in PLINK 1.90b3s or higher ^77^ for each cohort separately. QC steps on samples for KKNMS, KORA, and recMDD included removal of individuals with missing rate>2%, cryptic relatives (PI-HAT>0.0125), and genetic outliers (distance in first eight MDS components from mean >4 SD). QC steps on variants included removal of variants with a call rate<2%, MAF<1%, and HWE test *p*-value≤10^−6^. Furthermore, variants on non-autosomal chromosomes were excluded. QC steps for BoMa included removal of variants with call rate <5% (before sample removal), individuals with missing rate >2%, autosomal heterozygosity deviation (|F_het_|< 0.2), variants with call rate <5% (after sample removal), difference in variant missingness between cases and controls <2%, variants with MAF<1%, and variants with HWE test *p*-value≤10^−6^ in controls or *p*-value≤10^−10^ in cases. All non-autosomal chromosomes were excluded.

Imputation was also performed separately for each cohort using IMPUTE2 ^78^, following phasing in SHAPEIT ^79^, with the 1,000 genomes phase I reference panel (released in June 2014, ALL samples). QC of imputed probabilities was conducted in QCTOOL 1.4 (http://www.well.ox.ac.uk/~gav/qctool/). Imputed SNPs were excluded if MAF<1%, HWE test *p*-values ≤10^−6^ and the INFO metric <0.8. SNP coordinates are given according to hg19.

### Statistical analyses of genotype data (GWAS)

The GWAS for MDD was separately conducted on the two MDD cohorts, recMDD and BoMa. The GWAS for height was carried out using the population-based cohorts of KORA S3 and S4. PLINK 1.90b3s was used for these GWAS ^80^. Multidimensional scaling (MDS) was separately conducted on each GWAS cohort. For height the number of consecutive MDS components cumulatively explaining at least 80% of the genetic variance were selected as covariates (n=8) and for MDD the number of MDS components was selected based on Cattell’s scree test (n=3). Sex, age, and MDS components were used as covariates in logistic or linear regression. Data sets were combined using a fixed-effects meta analysis in METAL ^81^. The same covariates were retained for analysis with the DeepWAS method. For MS, the published GWAS results were used.

### Prediction of regulatory effects using DeepSEA

We employed DeepSEA ^36^ to determine the SNPs that potentially play an important role in human traits or diseases by acting through the alteration of regulatory elements. All the 919 DeepSEA chromatin features ^36^ representing regulatory information derived from the profiles of the ENCODE project ^16^ were considered as FUs. These units cover combinations of 201 different experimental annotations of epigenetically relevant information. This data includes 690 TF binding profiles for 160 different TFs, 125 DNase I hypersensitive site (DHS) profiles, and 104 histone mark profiles across 31 cell lines and 17 treatment conditions. The pretrained DeepSEA network (v0.94) was downloaded from http://deepsea.princeton.edu/help/ and the predictions and corresponding significance values of the regulatory effects, so-called e-values, of the set of all SNPs from three datasets were generated using a NVIDIA GeForce GTX TITAN X GPU (Maxwell). Significance was assessed by the method proposed by Zhou & Troyanskaya, which uses one million random SNPs from the 1,000 genomes project ^37^ as a background distribution to calculate e-values for each FU, by assessing the proportion of random variants with a bigger effect than that of observed variants. We applied an e-value cutoff of 5×10^−5^, to only take the SNPs associated with at least one FU into consideration (e.g., rs1035271 in GM12878:MEF2C). We refer to this set of SNPs as having a predicted regulatory effect.

### DeepWAS

#### Penalized regression models

Compared to the classical GWAS approaches where the trait of interest is regressed separately on each SNP, regularized regression approaches provide an alternative way to handle high dimensional data and to identify SNPs associated with the trait of interest using variable selection ^23^. Here we utilized L1-regularized logistic and linear regression (LASSO) with stability selection for variable selection implemented in the “stabs” R package ^82^. We decided to use LASSO regression instead of Elastic Net, as our regulatory SNPs were lowly correlated, i.e., of the 36,409 SNPs that were predicted to be regulatory for the imputed, high quality MS SNP set, 26,155 (72%) were uncorrelated (*r*^2^<0.2, window size of 50 SNPs, and a shift parameter of 5), with a mean *r*^2^ per FU ranging from 0.001 to 0.3. Furthermore, the fine-tuning of the Elastic Net meta-parameter L2 (alpha) is more difficult and time-consuming (for more details please see ^23^). Therefore, for each of 919 FUs, we fitted LASSO models via stability selection procedure in order to estimate stable statistical associations between the disease or trait status (binary for MS and MDD, continuous for height) and SNPs in the context of a specific cell line. Linear models fitted to datasets with a continuous response are described as follows:

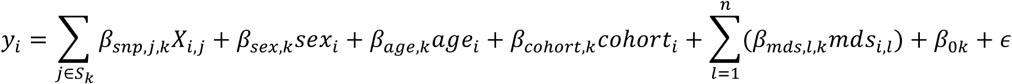

In each LASSO model, only the SNPs that significantly affect the binding of a specific chromatin feature in a specific cell line were included. This is represented in the equation by the summation over the elements of *S_k_*, which represents the set of SNPs that have an impact on FU *k*. Subscript *k* is used as a model index, which denotes a different model for each FU. We refer to the genotype matrix (e.g., 15,283 × 36,409 for the KKNMS dataset) as *X* where the rows are individuals (subscript *i*) and columns are SNPs (subscript *j*). Genotype in the *X* matrix is encoded using the dosage representation of each SNP in an additive model such that the final encoding of a SNP is *X_ij_* = 2×*P*(*AA_ij_*) + *P*(*Aa_ij_*) where *P*(*AA_ij_*) and *P*(*Aa_ij_*) are probabilities for being homozygous for the minor allele and heterozygous for individual *i* and SNP *j. X_ij_* is therefore a continuous value between 0 and 2. In addition to the SNP predictors, we used sex (binary), age (continuous), cohort dummy index, and MDS components (continuous) as covariates.

The model parameters (*β*) of the linear regression for continuous phenotype are optimized with L1 regularization, where the *λ* parameter represents the strength of the regularization. We fitted L1-regularized logistic and linear regression models (LASSO) using glmnet and stabs R packages ^83,84^. The regularization parameter *λ* is determined within the stability selection procedure, based on a provided probability cutoff and per-family error rate (PFER) values ^85^. Note that, unlike family-wise error rate and false discovery rate, which define a probability and an expected proportion, respectively, PFER defines the expected number of false positives and hence can be above 1.0. In total, n=100 subsample replicates were used for each model fit with a subsample size of ⌊n/2⌋. The probability cutoff represents how frequently a variable must be selected in LASSO models fitted to these replicates in order to be called a deepSNP. For more details of the relationship between the *λ* parameter, probability cutoff, and PFER values please see ^86^.

#### DeepWAS application

DeepWAS was conducted on the KKNMS dataset for MS, on the MDDC dataset for MDD, and on the KORA dataset for height. Sex, age, cohort membership, and significant MDS components were used as covariates in DeepWAS. A selection probability >0.7 and a PFER of 1.0 were used to identify significant trait associations (deepSNPs).

### Functional annotation of deepSNPs

#### DeepWAS (Encode) tissue category

Encode cell type information was download from https://genome.ucsc.edu/encode/cellTypes.html and tissue categories were extracted from the column „tissue”.

#### Roadmap *cis*-regulatory elements (ChromHMM)

*Cis*-regulatory elements identified by the Roadmap Epigenomics Project ^87^ were downloaded as segmentation files of core 15-state ChromHMM model for 111 epigenomes from the Roadmap epigenomics web portal (http://egg2.wustl.edu/roadmap/data/byFileType/chromhmmSegmentations/ChmmModels/coreMarks/jointModel/final/all.mnemonics.bedFiles.tgz) in BED format. ChromHMM used five core marks (H3K4me3, H3K4me1, H3K36me3, H3K27me3, H3K9me3) from each of the 111 reference epigenomes and learned a set 15-chromatin state definitions per genomic segment. We overlapped the DeepWAS SNPs with those chromatin states based on the exact genomic position. All roadmap epigenomes were grouped into broader tissue groups, which were used for mapping between DeepWAS and ChromHMM results.

#### Genomic region annotation

DeepWAS SNPs were overlapped with genomic annotation from UCSC for the hg19 genome build using “TxDb.Hsapiens.UCSC.hg19.knownGene” and “ChIPseeker” R packages ^88,89^.

#### Tissue enrichment with SNPsea and FANTOM5 CAGE data

SNPsea version 1.0.3 ^90^ was downloaded from https://github.com/slowkow/snpsea/. Tissue enrichment was performed with the command line interface of SNPsea. All data and annotation files were the default ones provided by SNPsea (tissue-specific gene expression from ~400 cell types), namely the *FANTOM2014.gct.gz* gene expression file was used with default *NCBIgenes2013.bed.gz, TGP2011.bed.gz, Lango2010.txt.gz* values for “gene-intervals”, “snp-intervals”, and “null-snps” options, respectively.

### DNA methylation data

For a subset of the recMDD cohort (n=166 MDD cases), genomic DNA was extracted from whole blood using the Gentra Puregene Blood Kit (QIAGEN). DNA quality and quantity of both was assessed with the NanoDrop 2000 Spectrophotometer (Thermo Scientific) and Quant-iT Picogreen (Invitrogen). Genomic DNA was bisulfite converted using the Zymo EZ-96 DNA Methylation Kit (Zymo Research) and DNA methylation levels were assessed for >480,000 CpG sites using the Illumina HumanMethylation450 BeadChips. Hybridization and processing were performed according to the manufacturer’s instructions. QC of methylation data, including intensity read outs, filtering (detection *p*-value > 0.01 in at least 75% of the samples), cellular composition estimation according to ^91^, as well as beta calculation were done using the minfi Bioconductor R package v1.18.2 ^92^. Filtered beta values were reduced by eliminating any CpG sites on sex chromosomes, as well as probes found to have SNPs at the CpG site itself or in the single-base extension site with a MAF ≥1% in the 1,000 genomes project EUR population and/or non-specific binding probes according to ^93^. Additionally, we performed a re-alignment of the array probe sequences using Bismark ^94^. This yielded a total of around 425,000 CpG sites for further analysis. The data were then normalized using functional normalization ^95^. Technical batch effects were identified by inspecting the association of the first principal components of the methylation levels with plate and plate position as technical batches. The data were then adjusted using ComBat ^96^. CpG coordinates are given according to hg19.

DNA methylation data was available for a subset of the KORA study (n= 1,802 F4 individuals). DNA methylation was measured with the Illumina HumanMethylation450K BeadChips. Sample preparation and measurement have been described previously ^97^. Intensity values were extracted from the idat files using the Bioconductor R package minfi, with subsequent background correction performed with lumi ^98^. Probes with a detection *p*-value > 0.01 or summarized by less than 3 functional beads were set to missing. A sample-wise call rate of 80% was applied, and color bias adjustment using smooth quantile normalization was performed. Finally, beta-mixture quantile normalization was performed on the probes (Bioconductor R package watermelon ^99^) to correct for the Inf I/Inf II distribution shift.

### Gene expression data

Gene expression analysis of the a subset of the KKNMS cohort (n=319 DE1 MS cases) was performed using llumina HT-12 v4 Expression BeadChips and published and described previously in ^27^. For a subset of the KORA cohort (n=1,002 F4 individuals) gene expression profiling was performed using the Illumina HT-12 v3 Expression BeadChips and described previously in ^100^.

### Statistical analysis of gene expression and methylation data

For the recMDD cohort, linear regression models were fit for each probe to test the relationship between the DNA methylation (beta values) and genotype (in dosage format) within 1Mb up- or downstream of the SNP using the R package MatrixEQTL ^101^. Sex, blood cell counts and genotype MDS components to correct for possible admixture effects were included as covariates. Significance after multiple testing was adjusted using a false discovery rate (FDR) of 5%.

For the KORA cohort, the OmicAbel software^102^ was used to test the association between methylation beta values and SNP dosage, adjusting for age, sex, and blood cell count. A total of 1,731 individuals had valid methylation and genetic data. Significance was defined after Bonferroni correction at a *p*-value of 1e-14. To examine the relationship between genetic variation and gene expression in KORA (n=711 individuals with valid genetic and expression data), we first derived residuals for gene transcript expression using linear regression of log_2_-transformed gene transcript levels against sex, age, RNA integrity number, RNA amplification plate, and sample storage time. Expression residuals were then used as outcome variables in a linear regression model with SNP dosage as the independent variable. Data analysis was performed using MatrixEQTL and significant meQTLs were filtered at an FDR of 5%.

For the analysis of association between genotype and expression levels in the KKNMS cohort, we used MatrixEQTL with sex, age, blood cell counts, and genotype MDS components as covariates. Significance after multiple testing was adjusted using an FDR of 5%.

Using published QTL data from BIOS whole peripheral blood DNA of 3,841 and mRNA of 2,116 healthy samples (eQTL, meQTL, eQTM), downloaded from http://genenetwork.nl/biosqtlbrowser, CMC eQTLs downloaded from https://www.synapse.org/#!Synapse:syn4622659, and GTEx eQTLs downloaded from https://gtexportal.org/home/datasets, we were able to intersect our deepSNPs and their LD proxies (*r*^2^≥0.5) with transcriptionally relevant data. The eQTLs obtained from GTEx were filtered on gene *p*<0.05 and eQTLs obtained from BIOS and CMC were filtered on FDR <0.05. We used permutation tests to determine if deepSNPs or their LD proxies were enriched in BIOS QTLs. For each phenotype, we compared the overlap of deepSNPs and BIOS QTL SNPs to the average overlap from 1,000 equally sized sets of regulatory SNPs and BIOS QTL SNPs.

#### QTL network

For the visualization of deepSNP-QTL interactions, we set up an Neo4j^103^ instance. The Neo4j community edition version 3.4.0 was downloaded from https://neo4j.com. All DeepWAS results were inserted to the database. The graph structure consisted of genes, transcription factors, CpGs, SNPs, cell lines, and tissues that are connected to each other through dummy nodes representing each deepSNP. Dummy nodes are especially important in cases where a deepSNP is predicted to be active in more than one FU, e.g. TF1:CL1 and TF2:CL2. In this case, connecting these 4 elements directly to the deepSNP leads to ambiguity about the FU, which can be misinterpreted as TF1:CL2 or TF2:CL1, since the information of the FU is lost. Using dummy nodes avoids this confusion by providing a SNP-FU link.

## Acknowledgements

We thank Michael Laimighofer and Ivan Kondofersky for feedback on the statistical approach. We would like to thank Richa Batra, Linda Krause, Christoph Orgis, Karolina Worf, Matthias Heinig and Martin Preusse for useful discussions on the approach.

## Ethics declaraion

All studies were approved by the local ethics committee and all individuals gave written informed consent. All experimental methods comply with the Helsinki Declaration.

## Author contributions

Conception and design: JA, GE and NSM; recruitment, acquisition and pre-processing of DE genotype and expression data: RG, BH, FL, SN, FP, HW, SHH, TK, ML, RR, TFMA and BMM; recruitment, acquisition and pre-processing of BoMa genotype data: MR, JF and JA; recruitment, acquisition and pre-processing of KORA genotype methylation and expression data: CG, TM, AP, KS, TFMA, MW and BK; recruitment, acquisition and pre-processing of recMDD genotype and methylation data: SL, BMM, JA, JM and SI; drafting or revising the manuscript: JA, GE, TFMA, EBB, TK, JF and NSM; analysis and interpretation of data: JA, GE, TFMA, EBB and NSM.

## Data availability

DeepWAS source code is available at https://zenodo.org/record/59282#.W82wEy-B2L4 through Zenodo. The informed consents given by KORA study participants do not cover data posting in public databases. However, data are available upon request from KORA-gen (https://epi.helmholtz-muenchen.de/). Data requests can be submitted online and are subject to approval by the KORA Board. KKNMS data are available upon approved request. MDDC genotype data are available upon approved request and DNAm data will be made available on GEO.

## References

1. Visscher, P. M., Brown, M. A., McCarthy, M. I. & Yang, J. Five Years of GWAS Discovery. The American Journal of Human Genetics 90, 7–24 (2012).

2. Edwards, S. L., Beesley, J., French, J. D. & Dunning, A. M. Beyond GWASs: illuminating the dark road from association to function. The American Journal of Human Genetics 93, 779–797 (2013).

3. Tak, Y. G. & Farnham, P. J. Making sense of GWAS: using epigenomics and genome engineering to understand the functional relevance of SNPs in non-coding regions of the human genome. Epigenetics & Chromatin 8, 57 (2015).

4. Maurano, M. T. et al. Systematic localization of common disease-associated variation in regulatory DNA. Science 337, 1190–1195 (2012).

5. Hindorff, L. A. et al. Potential etiologic and functional implications of genome-wide association loci for human diseases and traits. Proc Natl Acad Sci USA 106, 9362–9367 (2009).

6. Robertson, G. et al. Genome-wide profiles of STAT1 DNA association using chromatin immunoprecipitation and massively parallel sequencing. Nat Methods 4, 651–657 (2007).

7. Coetzee, S. G., Coetzee, G. A. & Hazelett, D. J. motifbreakR: an R/Bioconductor package for predicting variant effects at transcription factor binding sites: Fig. 1. Bioinformatics 10, btv470 (2015).

8. Thomas-Chollier, M. et al. Transcription factor binding predictions using TRAP for the analysis of ChIP-seq data and regulatory SNPs. Nat Protoc 6, 1860–1869 (2011).

9. Gamazon, E. R. et al. A gene-based association method for mapping traits using reference transcriptome data. Nat Genet 47, 1091–1098 (2015).

10. Gamazon, E. R. et al. Enrichment of cis-regulatory gene expression SNPs and methylation quantitative trait loci among bipolar disorder susceptibility variants. Molecular Psychiatry 18, 340–346 (2012).

11. Tehranchi, A. K. et al. Pooled ChIP-Seq Links Variation in Transcription Factor Binding to Complex Disease Risk. Cell 165, 730–741 (2016).

12. Kheradpour, P. et al. Systematic dissection of regulatory motifs in 2000 predicted human enhancers using a massively parallel reporter assay. Genome Research 23, 800–811 (2013).

13. Shen, S. Q. et al. Massively parallel cis-regulatory analysis in the mammalian central nervous system. Genome Research 26, 238–255 (2016).

14. Smith, R. P. et al. Massively parallel decoding of mammalian regulatory sequences supports a flexible organizational model. Nature Publishing Group 45, 1021–1028 (2013).

15. Zhou, J. & Troyanskaya, O. G. Predicting effects of noncoding variants with deep learning-based sequence model. Nat Meth 12, 931–934 (2015).

16. ENCODE Project Consortium et al. An integrated encyclopedia of DNA elements in the human genome. Nature 489, 57–74 (2012).

17. Roadmap Epigenomics Consortium et al. Integrative analysis of 111 reference human epigenomes. Nature 518, 317–330 (2015).

18. Zhou, J. et al. Deep learning sequence-based ab initio prediction of variant effects on expression and disease risk. Nature Publishing Group 1–13 (2018). doi:10.1038/s41588-018-0160-6

19. Zhou, J. et al. Whole-genome deep learning analysis reveals causal role of noncoding mutations in autism. (2018). doi:10.1101/319681

20. Sundaram, L. et al. Predicting the clinical impact of human mutation with deep neural networks. Nat Genet 50, 1161–1170 (2018).

21. Tibshirani, R. Regression shrinkage and selection via the lasso: a retrospective. Journal of the Royal Statistical Society: Series B (Statistical Methodology) 73, 273–282 (2011).

22. Thormann, V. et al. Genomic dissection of enhancers uncovers principles of combinatorial regulation and cell type-specific wiring of enhancer–promoter contacts. Nucleic Acids Res. 46, 2868–2882 (2018).

23. Waldmann, P., Mészáros, G., Gredler, B., Fuerst, C. & Sölkner, J. Evaluation of the lasso and the elastic net in genome-wide association studies. Front Genet 4, 270 (2013).

24. Ayers, K. L. & Cordell, H. J. SNP Selection in genome-wide and candidate gene studies via penalized logistic regression. Genet. Epidemiol. 34, 879–891 (2010).

25. Briollais, L. et al. A Bayesian graphical model for genome-wide association studies (GWAS). Ann. Appl. Stat. 10, 786–811 (2016).

26. Dehman, A., Ambroise, C. & Neuvial, P. Performance of a blockwise approach in variable selection using linkage disequilibrium information. BMC Bioinformatics 16, 148 (2015).

27. Andlauer, T. F. M. et al. Novel multiple sclerosis susceptibility loci implicated in epigenetic regulation. Science Advances 2, e1501678 (2016).

28. Muglia, P. et al. Genome-wide association study of recurrent major depressive disorder in two European case–control cohorts. Mol Psychiatry 15, 589–601 (2008).

29. Wichmann, H.-E., Gieger, C., Illig, T.MONICA/KORA Study Group. KORA-gen--resource for population genetics, controls and a broad spectrum of disease phenotypes. Gesundheitswesen 67 Suppl 1, S26–30 (2005).

30. Sullivan, P. F., Neale, M. C. & Kendler, K. S. Genetic Epidemiology of Major Depression: Review and Meta-Analysis. Am J Psychiatry 157, 1552–1562 (2000).

31. Patsopoulos, N. A. Genetics of Multiple Sclerosis: An Overview and New Directions. Cold Spring Harb Perspect Med 8, (2018).

32. Jelenkovic, A. et al. Genetic and environmental influences on height from infancy to early adulthood: An individual-based pooled analysis of 45 twin cohorts. Sci Rep 6, 246 (2016).

33. Lango-Allen, H. et al. Hundreds of variants clustered in genomic loci and biological pathways affect human height. Nature 467, 832–838 (2010).

34. Patsopoulos, N. A. Genetics of Multiple Sclerosis: An Overview and New Directions. Cold Spring Harb Perspect Med 8, a028951 (2018).

35. Wray, N. R. et al. Genome-wide association analyses identify 44 risk variants and refine the genetic architecture of major depression. Nat Genet 50, 668–681 (2018).

36. Zhou, J. & Troyanskaya, O. G. Predicting effects of noncoding variants with deep learning-based sequence model. Nat Methods 12, 931–934 (2015).

37. Consortium, 1. G. P. et al. A map of human genome variation from population-scale sequencing. Nature 467, 1061–1073 (2010).

38. Rietschel, M. et al. Genome-wide association-, replication-, and neuroimaging study implicates HOMER1 in the etiology of major depression. Biological Psychiatry 68, 578–585 (2010).

39. Holle, R., Happich, M., Löwel, H., Wichmann, H.-E. MONICA/KORA Study Group. KORA--a research platform for population based health research. Gesundheitswesen 67 Suppl 1, S19–25 (2005).

40. Aune, D. et al. Height and pancreatic cancer risk: a systematic review and meta-analysis of cohort studies. Cancer Causes Control 23, 1213–1222 (2012).

41. Ernst, J. & Kellis, M. Chromatin-state discovery and genome annotation with ChromHMM. Nat Protoc 12, 2478–2492 (2017).

42. Andersson, R. et al. An atlas of active enhancers across human cell types and tissues. Nature 507, 455–461 (2014).

43. - International Multiple Sclerosis Genetics Consorti et al. The Multiple Sclerosis Genomic Map: Role of peripheral immune cells and resident microglia in susceptibility. bioRxiv 1–43 (2017). doi:10.1101/143933

44. Bonder, M. J. et al. Disease variants alter transcription factor levels and methylation of their binding sites. (2015). doi:10.1101/033084

45. Zhernakova, D. V. et al. Identification of context-dependent expression quantitative trait loci in whole blood. Nat Genet 49, 139–145 (2016).

46. Fromer, M. et al. Gene expression elucidates functional impact of polygenic risk for schizophrenia. Nature Neuroscience 19, 1442–1453 (2016).

47. GTEx Consortium. Erratum: Genetic effects on gene expression across human tissues. Nature 553, 530–530 (2018).

48. Høglund, R. A. & Maghazachi, A. A. Multiple sclerosis and the role of immune cells. World J Exp Med 4, 27–37 (2014).

49. - International Multiple Sclerosis Genetics Consorti et al. The Multiple Sclerosis Genomic Map: Role of peripheral immune cells and resident microglia in susceptibility. (2017). doi:10.1101/143933

50. Gudbjartsson, D. F. et al. Many sequence variants affecting diversity of adult human height. Nat Genet 40, 609–615 (2008).

51. Olivier, J. D. A., Åkerud, H., Skalkidou, A., Kaihola, H. & Sundström-Poromaa, I. The effects of antenatal depression and antidepressant treatment on placental gene expression. Front Cell Neurosci 8, 465 (2014).

52. Puno, M. R. & Lima, C. D. Structural basis for MTR4-ZCCHC8 interactions that stimulate the MTR4 helicase in the nuclear exosome-targeting complex. Proceedings of the National Academy of Sciences 115, E5506–E5515 (2018).

53. Wang, X. et al. Discovery and validation of sub-threshold genome-wide association study loci using epigenomic signatures. eLife 5, (2016).

54. Adzhubei, I. A. et al. A method and server for predicting damaging missense mutations. Nat Meth 7, 248–249 (2010).

55. Ng, P. C. & Henikoff, S. SIFT: Predicting amino acid changes that affect protein function. Nucleic Acids Res. 31, 3812–3814 (2003).

56. Visscher, P. M. et al. 10 Years of GWAS Discovery: Biology, Function, and Translation. Am. J. Hum. Genet. 101, 5–22 (2017).

57. O’Brien, J. S. & Kishimoto, Y. Saposin proteins: structure, function, and role in human lysosomal storage disorders. FASEB J. 5, 301–308 (1991).

58. Halmer, R., Walter, S. & Faßbender, K. Sphingolipids: important players in multiple sclerosis. Cell. Physiol. Biochem. 34, 111–118 (2014).

59. Chen, S. X. et al. The transcription factor MEF2 directs developmental visually driven functional and structural metaplasticity. Cell 151, 41–55 (2012).

60. Barbosa, A. C. et al. MEF2C, a transcription factor that facilitates learning and memory by negative regulation of synapse numbers and function. Proc Natl Acad Sci USA 105, 9391–9396 (2008).

61. Li, S., Alvarez, R. V., Sharan, R., Landsman, D. & Ovcharenko, I. Quantifying deleterious effects of regulatory variants. Nucleic Acids Res. 45, 2307–2317 (2017).

62. Kleiman, E., Jia, H., Loguercio, S., Su, A. I. & Feeney, A. J. YY1 plays an essential role at all stages of B-cell differentiation. Proceedings of the National Academy of Sciences 113, E3911–20 (2016).

63. Balkhi, M. Y., Wittmann, G., Xiong, F. & Junghans, R. P. YY1 Upregulates Checkpoint Receptors and Downregulates Type I Cytokines in Exhausted, Chronically Stimulated Human T Cells. iScience 2, 105–122 (2018).

64. Chihara, N. Dysregulated T cells in multiple sclerosis. Clin Exp Neuroimmunol 9, 20–29 (2018).

65. M, S. Stage-Specific Immune Dysregulation in Multiple Sclerosis. Journal of Interferon & Cytokine Research 34, 633–640 (2014).

66. Suhre, K. et al. Connecting genetic risk to disease end points through the human blood plasma proteome. Nature Communications 8, 14357 (2017).

67. Arloth, J. et al. Genetic Differences in the Immediate Transcriptome Response to Stress Predict Risk-Related Brain Function and Psychiatric Disorders. Neuron 86, 1189–1202 (2015).

68. Wohleb, E. S., Franklin, T., Iwata, M. & Duman, R. S. Integrating neuroimmune systems in the neurobiology of depression. Nat Rev Neurosci 17, 497–511 (2016).

69. Pruim, R. J. et al. LocusZoom: regional visualization of genome-wide association scan results. Bioinformatics 26, 2336–2337 (2010).

70. Rietschel, M. et al. Genome-wide association-, replication- and neuroimaging study implicates HOMER1 in the aetiology of major depression. Journal of Affective disorders 122, S38–S39 (2010).

71. Schmermund, A. et al. Assessment of clinically silent atherosclerotic disease and established and novel risk factors for predicting myocardial infarction and cardiac death in healthy middle-aged subjects: rationale and design of the Heinz Nixdorf RECALL Study. Risk Factors, Evaluation of Coronary Calcium and Lifestyle. Am. Heart J. 144, 212–218 (2002).

72. Völzke, H. et al. Cohort profile: the study of health in Pomerania. Int J Epidemiol 40, 294–307 (2011).

73. Berger, K. DHS. Bundesgesundheitsblatt - Gesundheitsforschung - Gesundheitsschutz 55, 816–821 (2012).

74. Müller, N. et al. IL-6 blockade by monoclonal antibodies inhibits apolipoprotein (a) expression and lipoprotein (a) synthesis in humans. Journal of Lipid Research 56, 1034–1042 (2015).

75. Krawczak, M. et al. PopGen: population-based recruitment of patients and controls for the analysis of complex genotype-phenotype relationships. Community Genet 9, 55–61 (2006).

76. Kohli, M. A. et al. The neuronal transporter gene SLC6A15 confers risk to major depression. Neuron 70, 252–265 (2011).

77. Chang, C. C. et al. Second-generation PLINK: rising to the challenge of larger and richer datasets. Gigascience 4, 7 (2015).

78. Howie, B. N., Donnelly, P. & Marchini, J. A flexible and accurate genotype imputation method for the next generation of genome-wide association studies. PLoS Genet 5, e1000529 (2009).

79. Delaneau, O., Zagury, J.-F. & Marchini, J. Improved whole-chromosome phasing for disease and population genetic studies. Nat Methods 10, 5–6 (2013).

80. Chang, C. C. et al. Second-generation PLINK: rising to the challenge of larger and richer datasets. Gigascience 4, 7 (2015).

81. Willer, C. J., Li, Y. & Abecasis, G. R. METAL: fast and efficient meta-analysis of genomewide association scans. Bioinformatics 26, 2190–2191 (2010).

82. Hofner, B., Boccuto, L. & Göker, M. Controlling false discoveries in high-dimensional situations: boosting with stability selection. BMC Bioinformatics 16, 144 (2015).

83. Simon, N., Friedman, J., Hastie, T. & Tibshirani, R. Regularization Paths for Cox’s Proportional Hazards Model via Coordinate Descent. J. Stat. Soft. 39, (2011).

84. Hofner, B., Boccuto, L. & Göker, M. Controlling false discoveries in high-dimensional situations: boosting with stability selection. BMC Bioinformatics 16, 144 (2015).

85. Dudoit, S., Shaffer, J. P. & Block, J. C. Multiple Hypothesis Testing in Microarray Experiments. Statistical Science 18, 71–103 (2003).

86. Meinshausen, N., Meier, L. & Bühlmann, P. p-Values for High-Dimensional Regression. Journal of the American Statistical Association 104, 1671–1681 (2009).

87. Roadmap Epigenomics Consortium et al. Integrative analysis of 111 reference human epigenomes. Nature 518, 317–330 (2015).

88. Carlson M, Maintainer BP (2015). TxDb.Hsapiens.UCSC.hg19.knownGene: Annotation package for TxDb.

89. Yu, G., Wang, L.-G. & He, Q.-Y. ChIPseeker: an R/Bioconductor package for ChIP peak annotation, comparison and visualization. Bioinformatics 31, 2382–2383 (2015).

90. Slowikowski, K., Hu, X. & Raychaudhuri, S. SNPsea: an algorithm to identify cell types, tissues and pathways affected by risk loci. Bioinformatics 30, 2496–2497 (2014).

91. Houseman, E. A. et al. DNA methylation arrays as surrogate measures of cell mixture distribution. BMC Bioinformatics 13, 86 (2012).

92. Aryee, M. J. et al. Minfi: a flexible and comprehensive Bioconductor package for the analysis of Infinium DNA methylation microarrays. Bioinformatics 30, 1363–1369 (2014).

93. Chen, Y.-A. et al. Discovery of cross-reactive probes and polymorphic CpGs in the Illumina Infinium HumanMethylation450 microarray. Epigenetics 8, 203–209 (2013).

94. Krueger, F. & Andrews, S. R. Bismark: a flexible aligner and methylation caller for Bisulfite-Seq applications. Bioinformatics 27, 1571–1572 (2011).

95. Fortin, J.-P. et al. Functional normalization of 450k methylation array data improves replication in large cancer studies. Genome Biol 15, 503 (2014).

96. Johnson, W. E. & Cheng, L. Adjusting batch effects in microarray expression data using empirical Bayes methods. Biostatistics 8, 118–127 (2007).

97. Zeilinger, S. et al. Tobacco smoking leads to extensive genome-wide changes in DNA methylation. PLoS ONE 8, e63812 (2013).

98. Du, P., Kibbe, W. A. & Lin, S. M. lumi: a pipeline for processing Illumina microarray. Bioinformatics 24, 1547–1548 (2008).

99. Pidsley, R. et al. A data-driven approach to preprocessing Illumina 450K methylation array data. BMC Genomics 14, 293 (2013).

100. Schurmann, C. et al. Analyzing illumina gene expression microarray data from different tissues: methodological aspects of data analysis in the metaxpress consortium. PLoS ONE 7, e50938 (2012).

101. Shabalin, A. A. Matrix eQTL: ultra fast eQTL analysis via large matrix operations. Bioinformatics 28, 1353–1358 (2012).

102. Fabregat-Traver, D. et al. High-Performance Mixed Models Based Genome-Wide Association Analysis with omicABEL software. F1000Res 3, 200 (2014).

103. Neo4J Developers, 2012. Neo4j. Graph NoSQL Database. Available at: (Accessed: 17 October 2018)

