## Supplementary material for "DeepWAS: Multivariate genotype-phenotype associations by directly integrating regulatory information using deep learning"

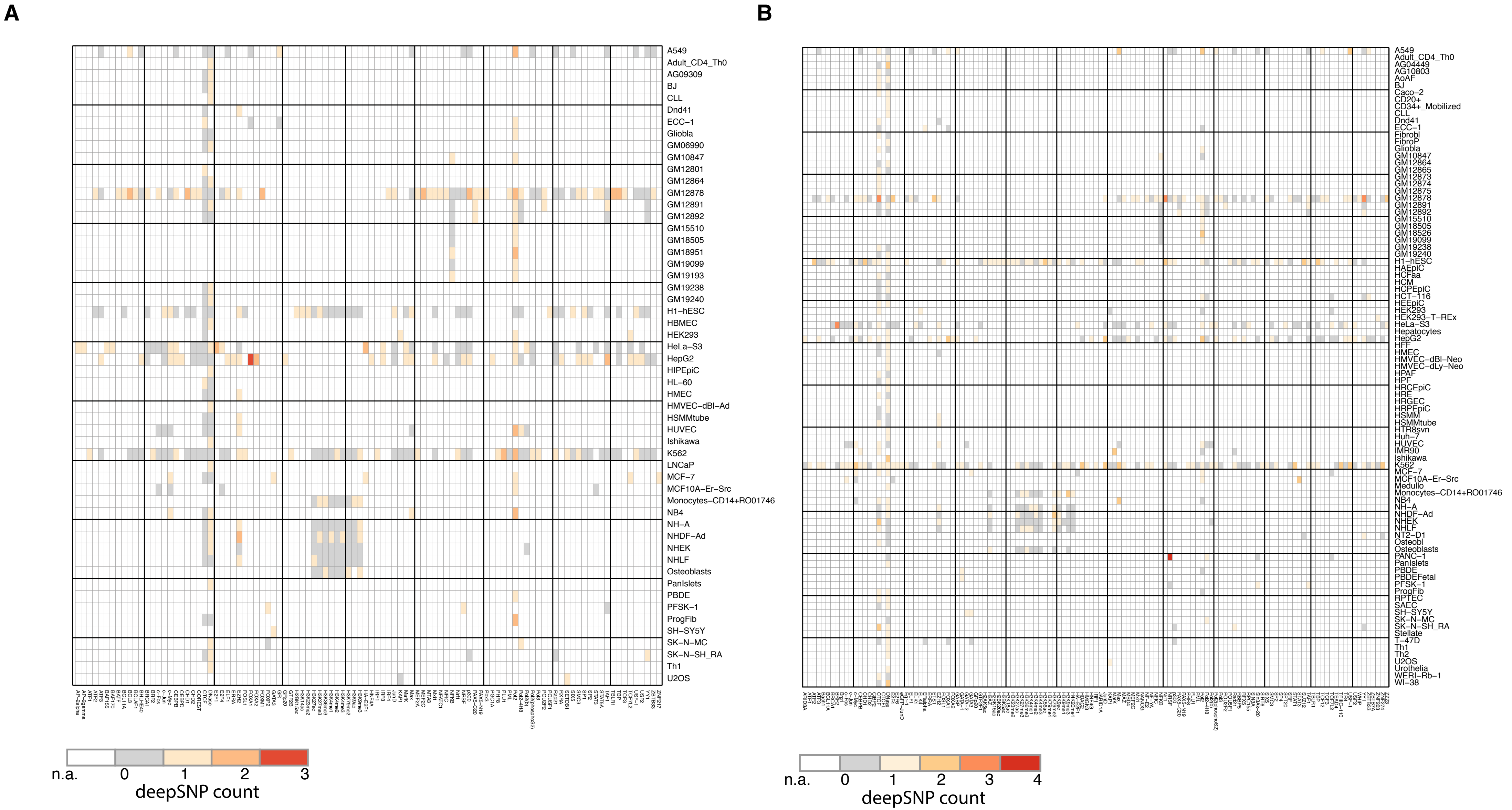


**Supplementary Figure 1**

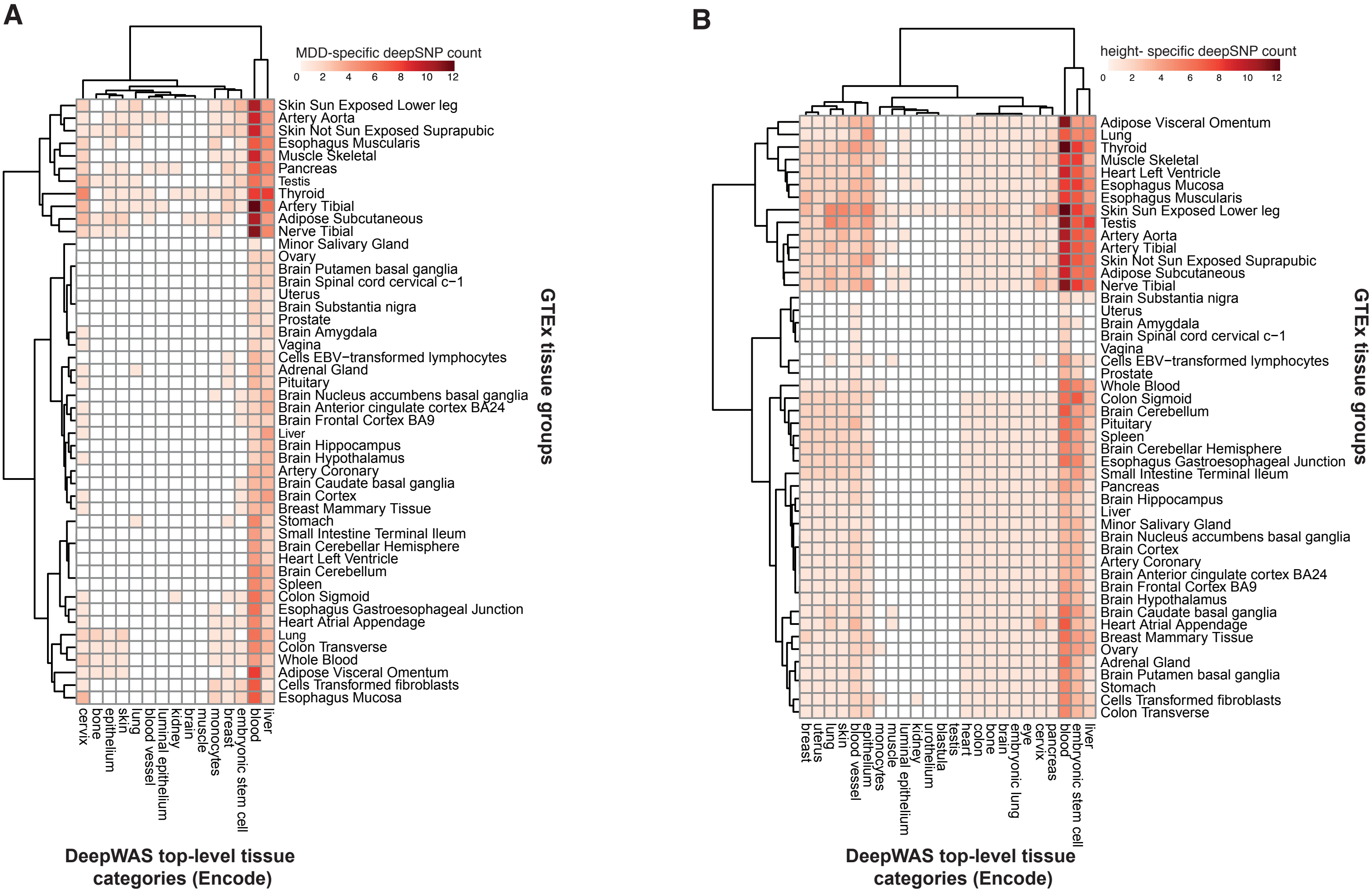


**Supplementary Figure 2**

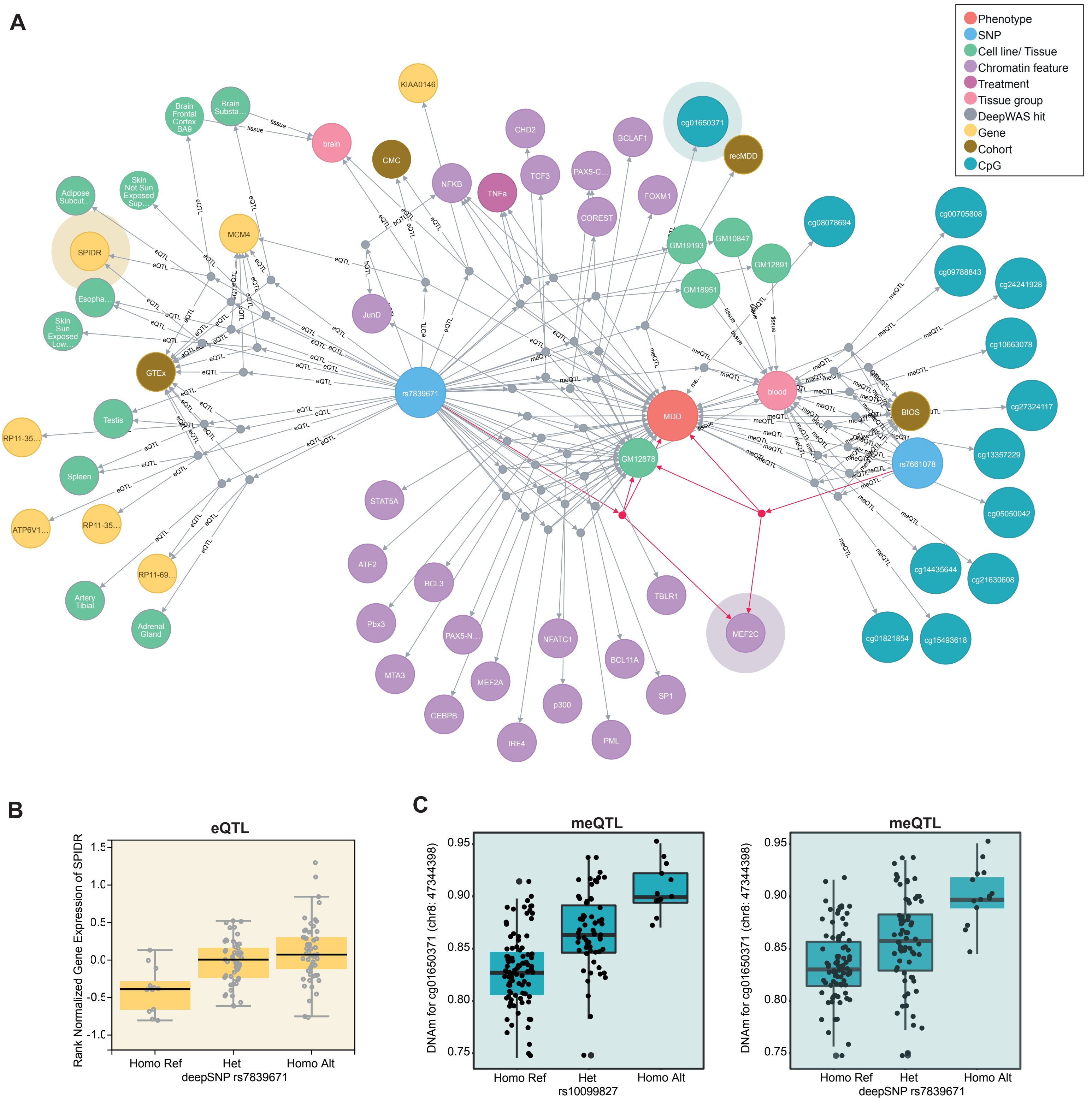


Supplementary Figure 3

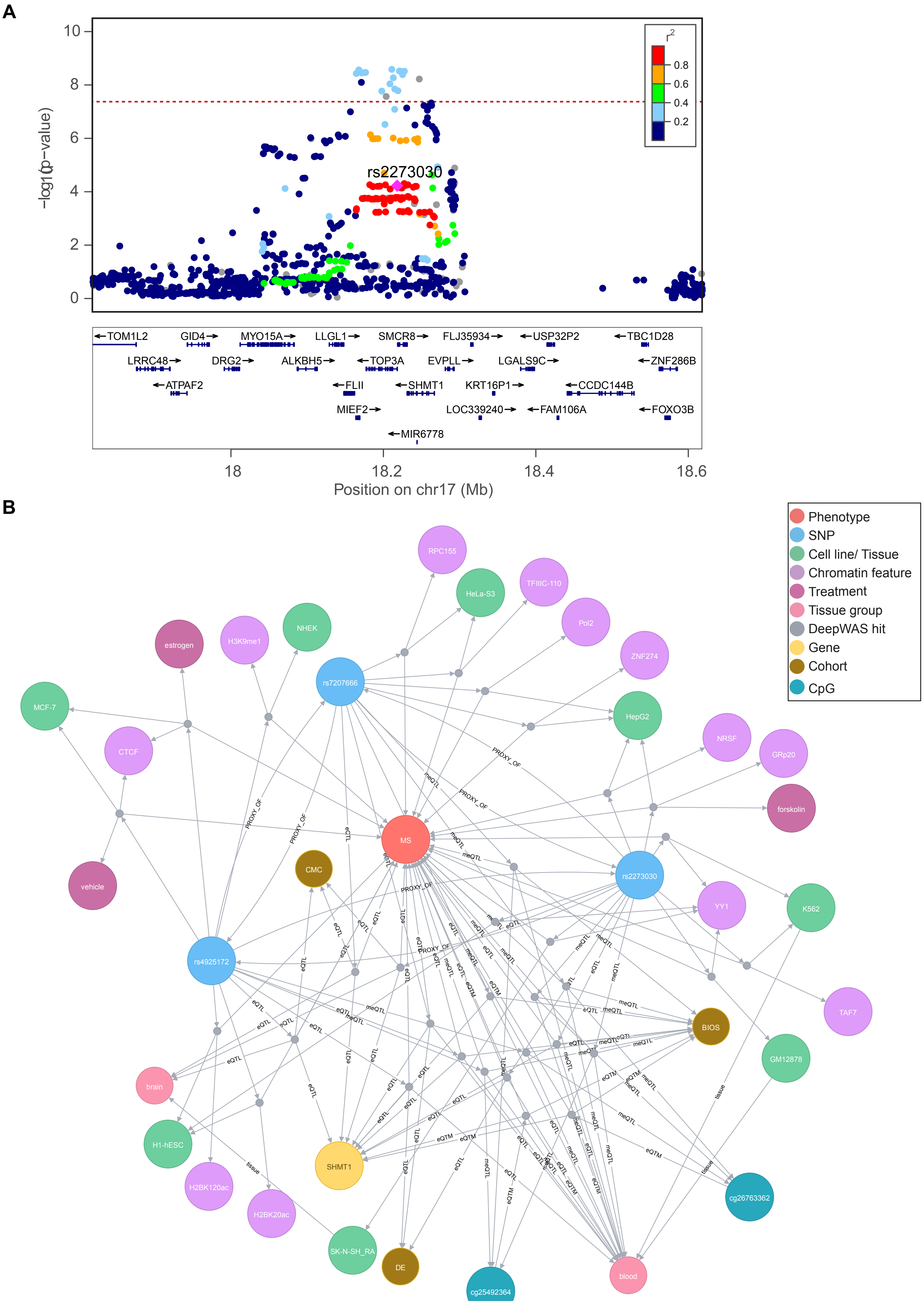


Supplementary Figure 4

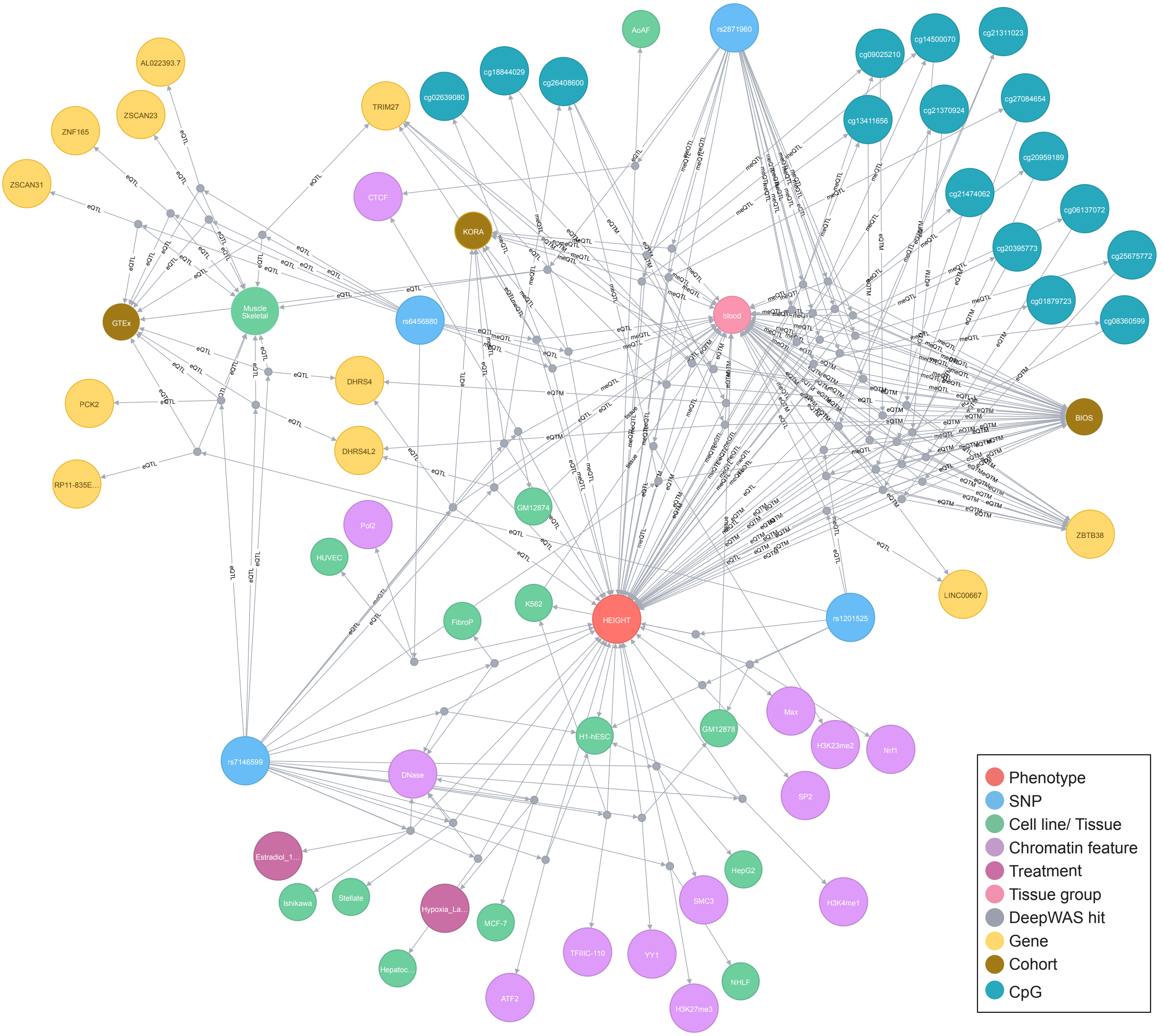


**Supplementary Figure 5**

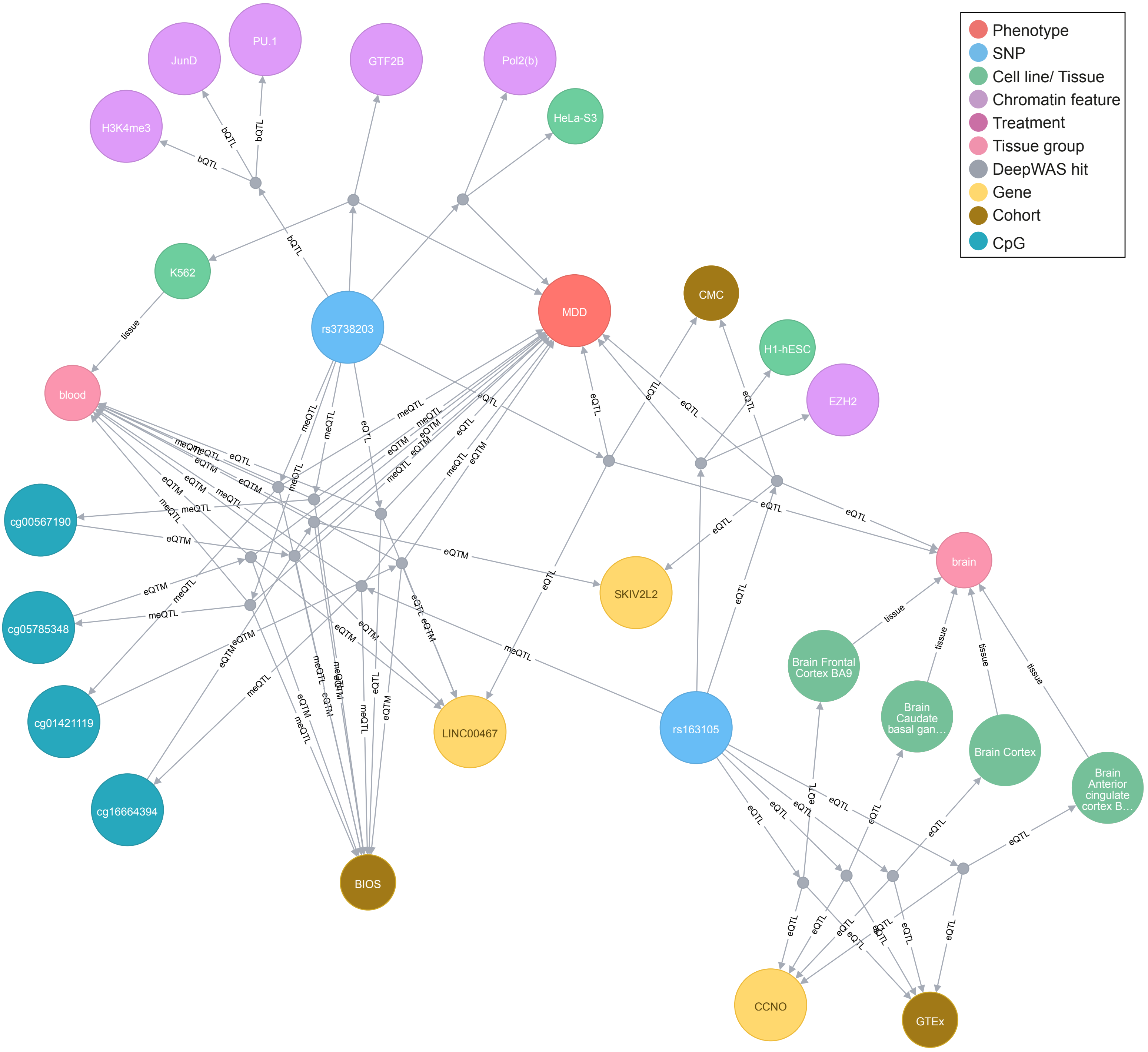


**Supplementary Figure 6**

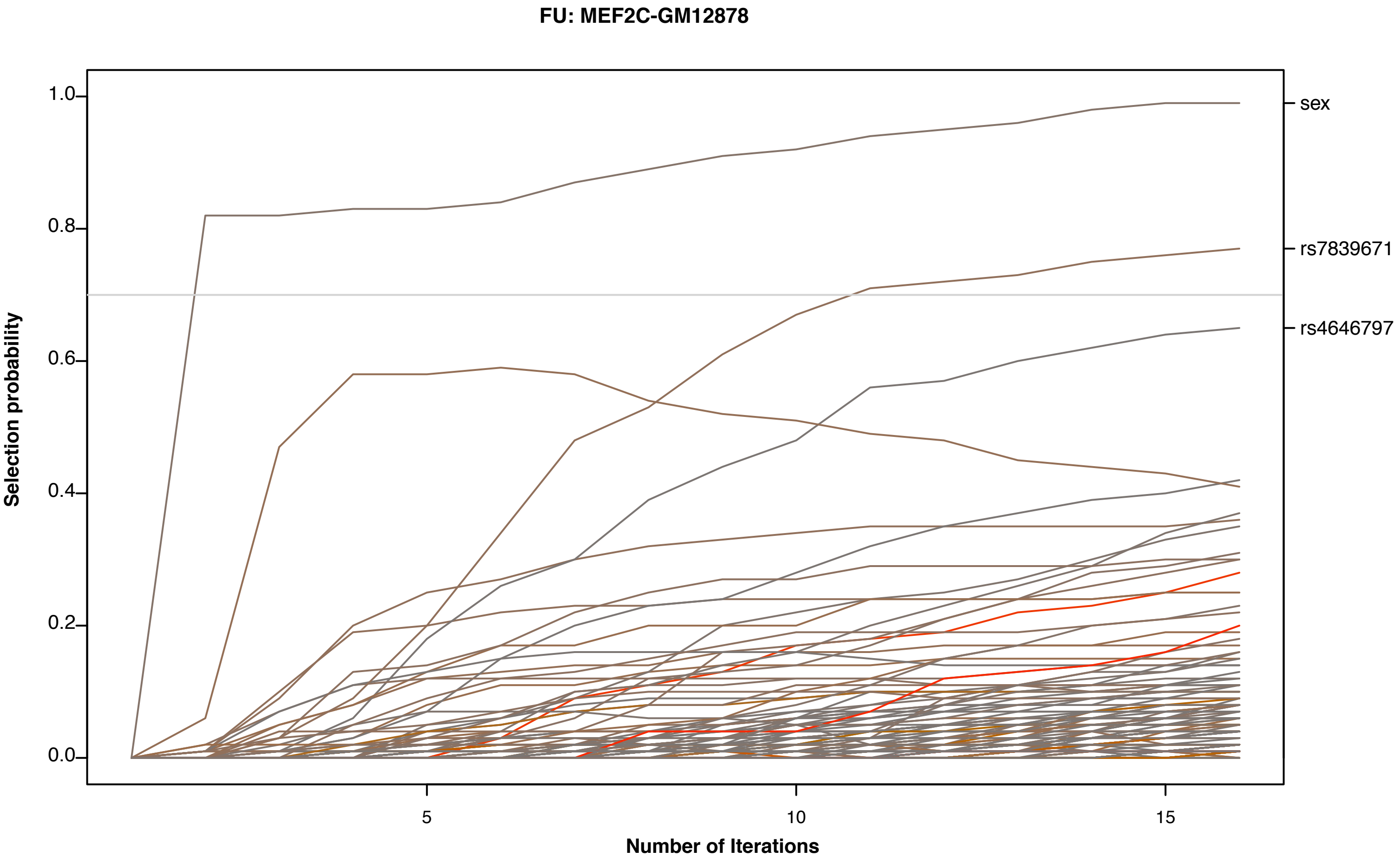


**Supplementary Figure 7**
